# ROLE OF FORELIMB MORPHOLOGY IN MUSCLE SENSORIMOTOR FUNCTIONS DURING LOCOMOTION IN THE CAT

**DOI:** 10.1101/2024.07.11.603106

**Authors:** Seyed Mohammadali Rahmati, Alexander N. Klishko, Ramaldo S. Martin, Nate E. Bunderson, Jeswin A. Meslie, T. Richard Nichols, Ilya A. Rybak, Alain Frigon, Thomas J. Burkholder, Boris I. Prilutsky

## Abstract

Previous studies established strong links between morphological characteristics of mammalian hindlimb muscles and their sensorimotor functions during locomotion. Less is known about the role of forelimb morphology in motor outputs and generation of sensory signals. Here, we measured morphological characteristics of 46 forelimb muscles from 6 cats. These characteristics included muscle attachments, physiological cross-sectional area (PCSA), fascicle length, etc. We also recorded full-body mechanics and EMG activity of forelimb muscles during level overground and treadmill locomotion in 7 and 16 adult cats of either sex, respectively. We computed forelimb muscle forces along with force- and length-dependent sensory signals mapped onto corresponding cervical spinal segments. We found that patterns of computed muscle forces and afferent activities were strongly affected by the muscle’s moment arm, PCSA, and fascicle length. Morphology of the shoulder muscles suggests distinct roles of the forelimbs in lateral force production and movements. Patterns of length-dependent sensory activity of muscles with long fibers (brachioradialis, extensor carpi radialis) closely matched patterns of overall forelimb length, whereas the activity pattern of biceps brachii matched forelimb orientation. We conclude that cat forelimb muscle morphology contributes substantially to locomotor function, particularly to control lateral stability and turning, rather than propulsion.

## INTRODUCTION

Quadrupedal mammals evolved their musculoskeletal and neural sensorimotor systems to perform various locomotor behaviors, such as searching for food, escaping predators, finding mates to produce offspring, etc. Evolutionary history and functional demands are critical factors defining the morphology of animal limbs and their sensorimotor functions (Shubin et al., 1997; Diogo and Molnar, 2014; Rothier et al., 2023). Mammalian forelimbs and hindlimbs have evolved many common morphological features. For instance, three-segment kinematic chains (i.e. upper arm-forearm-metacarpus and thigh-shank-metatarsus) are designed to save energy and improve stability (Kuznetsov, 1995; Fischer and Blickhan, 2006). A distal-to-proximal gradient of limb inertia with larger muscles located around proximal joints allows for faster and more economical swing and reaching movements (Gambaryan, 1974; Martin et al., 2010; Hudson et al., 2011a, b; Mathewson et al., 2012; Charles et al., 2016a; Kilbourne and Carrier, 2016) and paw shake responses (Prilutsky et al., 2022). A significant redundancy of muscle action, with over 30 muscles serving just 7 major degrees of freedom (Prilutsky and Zatsiorsky, 2002; Burkholder and Nichols, 2004; Bunderson et al., 2010; Martin et al., 2010; Ramalingasetty et al., 2021; Stark et al., 2021) is only partially resolved by considering non-sagittal actions. The presence of two-joint muscles in the hindlimb and forelimb contributes to regulating limb stiffness and impedance, inter-joint coordination, reducing exergy expenditure and muscle fatigue, and distribution of mechanical energy along the limb (Elftman, 1940; Nichols, 1994; Van Ingen Schenau et al., 1994; Prilutsky, 2000a, b). Similarities in limb morphology between the fore- and hindlimb correspond to a generally similar organization of excitatory and inhibitory somatosensory pathways. For example, the monosynaptic Ia muscle length-dependent pathways in the cat hindlimb (Eccles et al., 1957a; Eccles and Lundberg, 1958; Edgley et al., 1986; Nichols, 1994, 1999) and forelimb (Fritz et al., 1989; Illert, 1996; Caicoya et al., 1999), along with Ib pathways (Eccles et al., 1957b; Nichols, 2018) and recurrent inhibition pathways (Eccles et al., 1961; Illert and Wietelmann, 1989; Turkin et al., 1998) coordinate synergistic activities of agonists and antagonists at and across joints, although these pathways appear more complex in the forelimb (Fritz et al., 1989). The greater volume (the product of the mean physiological cross-sectional area and muscle fascicle length) of proximal versus distal muscles in the forelimbs and hindlimbs typically results in longer muscle moment arms of proximal muscles (Graham and Scott, 2003; Burkholder and Nichols, 2004; Hudson et al., 2011b, a; Charles et al., 2016b; Ramalingasetty et al., 2021). This translates in a higher acuity of length-dependent afferents to angle changes in proximal joints compared to distal ones (Hall and McCloskey, 1983; Burkholder and Nichols, 2000; Oh and Prilutsky, 2022). This feature of hip muscles could explain their important role in triggering the stance-swing (Kriellaars et al., 1994; Lam and Pearson, 2001) and swing-stance (McVea et al., 2005; Gregor et al., 2006) transitions during cat locomotion.

There are also differences in morphology and function between the forelimbs and hindlimbs. One of the most noticeable is the opposite flexion-extension directions in the homologous elbow and knee joints, which is likely caused by a much broader functional repertoire of the forelimbs, with unique roles in exploratory, hunting, reaching and grasping behaviors. Mass and volume of hindlimb muscles are greater than those of the forelimbs (Gambaryan, 1974; Hudson et al., 2011a, b; Ramalingasetty et al., 2021), although the forelimbs support a greater fraction of body weight than the hindlimbs due to a more rostral position of the body’s center of mass (Gray, 1944; Frigon et al., 2021). Larger muscle volume in the hindlimbs permits greater production of positive power/work and body acceleration compared to the forelimbs during locomotion (Pandy et al., 1988; Gregersen et al., 1998; Dutto et al., 2006; Jarrell et al., 2018). The forelimbs contribute more to body deceleration and the absorption of body energy (the production of negative power/work) than the hindlimbs during locomotion with constant speeds (Dutto et al., 2004; Dutto et al., 2006; Corbee et al., 2014; Farrell et al., 2014). It is expected that differences in morphology and mechanical functions between mammalian hindlimbs and forelimbs are also reflected in differences in their neural control.

Knowledge of limb morphology and its roles in sensorimotor functions is critically important for understanding the neural control of quadrupedal locomotion because the nervous system evolved in parallel with the musculoskeletal system, integrating feedforward central mechanisms in the brain and spinal cord with motion-dependent somatosensory feedback (Yakovenko and Drew, 2015; Danner et al., 2017; Grillner and El Manira, 2020; Frigon et al., 2021; Arber and Costa, 2022; Leiras et al., 2022; Dougherty, 2023). Previous studies established strong links between morphological characteristics of hindlimb muscles and their motor and sensory locomotor functions, especially in the cat (Spector et al., 1980; He et al., 1991; Nichols, 1994; Prochazka, 1999; Burkholder and Nichols, 2000; Burkholder and Lieber, 2001; Yakovenko et al., 2002; Ekeberg and Pearson, 2005; Bunderson et al., 2010; Charles et al., 2016a). However, little is known about the role of forelimb muscle morphology in producing motor outputs and generating length- and force-dependent sensory signals. This information is needed to understand contributions of the forelimbs in controlling quadrupedal locomotion.

Therefore, the goal of this study was to determine morphological characteristics of cat forelimb muscles and evaluate their contribution to locomotor kinetics and motion-dependent sensory feedback. We constructed a musculoskeletal model of the cat forelimb and validated select distal moment arms. We used kinematics and kinetics collected from walking cats to estimate muscle activations during walking. We validated these predictions against selected EMG recordings. Finally, we estimated proprioceptive afferent activities and mapped them onto corresponding cervical spinal segments. This information provided unprecedented insight into the role of mammalian forelimb morphology in motor and somatosensory functions during locomotion.

## METHODS

### Specimens and experimental subjects

This project involved two groups of animals. Fourteen forelimbs were harvested from 10 cats that had been euthanized following unrelated experiments involving only the hindlimb. Of these limbs, six were used for muscle architecture, of which four were also used to collect passive kinematics for joint center estimation, and the final eight limbs were used for moment arm measurements. Segment lengths of the six architecture forelimbs are summarized in Table 1. In vivo experiments described in this study were in agreement with US Public Health Service Policy on Humane Care and Use of Laboratory Animals. All experimental procedures were reviewed and approved by the Institutional Animal Care and Use Committees of Georgia Institute of Technology and Université de Sherbrooke. Our studies follow the ARRIVE 2.0 guidelines for animal studies (Percie du Sert et al., 2020). To minimize the number of animals, the same animals have been used in several studies to address different scientific questions. Here we report new experimental results obtained in cats that also participated in previous studies (Maas et al., 2010; Klishko et al., 2014; Farrell et al., 2015; Gregor et al., 2018; Klishko et al., 2021; Lecomte et al., 2022; Merlet et al., 2022; Audet et al., 2023; Lecomte et al., 2023; Mari et al., 2023; Park et al., 2023; Harnie et al., 2024; Mari et al., 2024).

**Table 1.** Segment lengths of forelimbs from which morphological characteristics were obtained(cm)

| Cat | Scapula | Humerus | Radius | Carpals+Metacarpals |
| --- | --- | --- | --- | --- |
| Cat A | 6.65 | 9.49 | 8.84 | 3.09 |
| Cat B | 7.26 | 10.39 | 9.30 | 3.59 |
| Cat C | 7.22 | 10.16 | 9.35 | 3.23 |
| Cat D | 7.28 | 10.05 | 9.06 | 3.27 |
| Cat F | 7.31 | 10.07 | 9.40 | 3.53 |
| Cat E | 7.04 | 9.56 | 9.07 | 3.29 |
| Mean±SD | 7.13±0.25 | 9.95±0.36 | 9.17±0.22 | 3.33±0.19 |

Forelimb kinematics and ground reaction forces during overground walking at self-selected speeds were recorded in a group of 7 adult female cats (1-2 years, mass 3.20 ± 0.34 kg; Tables 2 and 3). Electromyographic activity (EMG) of forelimb muscles during treadmill walking at a speed of 0.4 m/s were recorded in a separate group of 16 adult female and male cats (1-2 years, mass 4.82 ± 0.99 kg, Table 4).

**Table 2.** Characteristics of cats participated in motion capture experiments.

| Cat | Sex | Mass,<br>kg | Scapula<br>length,<br>cm | Upper<br>arm<br>length,<br>cm | Forearm<br>length,<br>cm | Carpals+<br>Metacarpals<br>length, cm | Number<br>of<br>analyzed<br>cycles | Cycle<br>duration<br>(mean±<br>SD), s | Duty<br>cycle<br>(mean±<br>SD) |
| --- | --- | --- | --- | --- | --- | --- | --- | --- | --- |
| BL | F | 3.00 | 6.50 | 8.75 | 8.90 | 3.4 | 35 | 0.695±<br>0.026 | 0.63±<br>0.01 |
| IN | F | 3.35 | 5.05 | 9.09 | 9.66 | 2.3 | 29 | 0.688±<br>0.030 | 0.65±<br>0.01 |
| KA | F | 3.15 | 5.30 | 9.55 | 8.91 | 3.0 | 13 | 0.634±<br>0.024 | 0.64±<br>0.01 |
| MA | F | 3.05 | 6.20 | 8.95 | 10.00 | 2.4 | 44 | 0.582±<br>0.021 | 0.63±<br>0.01 |
| RY | F | 2.80 | 5.75 | 10.00 | 10.00 | 2.05 | 19 | 0.867±<br>0.025 | 0.64±<br>0.01 |
| SV | F | 3.86 | 5.75 | 9.49 | 9.49 | 3.0 | 68 | 0.619±0.<br>003 | 0.65±<br>0.002 |
| TU | F | 3.20 | 5.85 | 9.10 | 9.30 | 2.15 | 3 | 0.747±<br>0.032 | 0.62±<br>0.02 |
| Mean±<br>SD | - | 3.20±<br>0.34 | 5.77±<br>0.49 | 9.28±<br>0.43 | 9.47±<br>0.46 | 2.61±<br>0.52 | - | 0.692±<br>0.093 | 0.64±<br>0.01 |

Note: Measurements of segment lengths were performed on sedated cats by an anthropometer.
The number of cycles analyzed corresponds to the maximum number of cycles analyzed for a selected mechanical variable.

**Table 3.** Inertial parameters of forelimb segments of 7 cats for whom joint moments of force were computed (mean ± SD)

| Parameter | Scapula | Upper arm | Forearm | Carpals | Digits |
| --- | --- | --- | --- | --- | --- |
| Center of mass location, cm | 2.75±0.24 | 4.53±0.21 | 4.30±2.1 | 1.37±0.27 | 1.27±0.41 |
| Mass, g | 63.1±4.0 | 81.0±6.6 | 41.6±3.3 | 7.5±1.6 | 5.1±1.3 |
| Moment of inertia, g · mm<sup>2</sup> | 35004±2833 | 59760±9617 | 35388±4338 | 587±179 | - |

Notes. Segment inertial parameters were computed from cat mass and forelimb segment length using regression equations developed in (Hoy and Zernicke, 1985). Center of mass location is determined from the proximal segment end. Moment of inertia is determined with respect to the medio-lateral axis at the segment center of mass. Digits are considered a point mass, i.e., there is no moment of inertia with respect to the digits center of mass.

**Table 4.** Characteristics of cats participated in EMG recordings of forelimb muscles.

| Cat | Mass, kg | Sex | Number of<br>analyzed<br>cycles | Cycle duration<br>(mean±SD), s | Duty cycle<br>(mean±SD) |
| --- | --- | --- | --- | --- | --- |
| BE | 5.10 | Male | 123 | 0.896±0.118 | 0.70±0.05 |
| DE | 4.65 | Male | 14 | 1.183±0.058 | 0.74±0.03 |
| GR | 4.80 | Female | 17 | 1.030±0.107 | 0.71±0.05 |
| HO | 5.90 | Male | 17 | 1.059±0.089 | 0.77±0.02 |
| JA | 6.10 | Male | 10 | 0.975±0.179 | 0.72±0.06 |
| KA | 4.00 | Female | 34 | 0.930±0.056 | 0.71±0.02 |
| KI | 5.60 | Male | 25 | 1.046±0.120 | 0.72±0.04 |
| LIM | 3.55 | Female | 54 | 0.908±0.079 | 0.74±0.03 |
| LIS | 3.15 | Female | 18 | 1.028±0.198 | 0.69±0.06 |
| MO | 5.34 | Male | 61 | 1.083±0.116 | 0.69±0.05 |
| OS | 6.30 | Male | 14 | 0.971±0.043 | 0.74±0.02 |
| PA | 3.99 | Female | 31 | 0.822±0.078 | 0.70±0.05 |
| RI | 4.80 | Male | 48 | 1.185±0.185 | 0.75±0.04 |
| TOK | 3.95 | Female | 31 | 1.092±0.223 | 0.72±0.08 |
| TOR | 6.00 | Male | 18 | 1.079±0.127 | 0.75±0.03 |
| VE | 3.87 | Female | 16 | 1.106±0.158 | 0.74±0.05 |
| Mean±SD | 4.82±0.99 | - | - | 1.025±0.102 | 0.72±0.02 |

### Mathematical Model

#### Strategy

The model includes 26 intrinsic muscles, subdivided to account for 40 tendons of insertion, with forces estimated by Hill-style models (Bunderson et al., 2010; Prilutsky et al., 2016) and rigid tendons, and afferent feedback estimated by Prochazka-style spindle and tendon organ models (Prochazka, 1999). Joint centers were defined by motion-captured passive mechanics and further refined to match measured wrist and elbow moment arms. Muscle attachment points were measured using a coordinate measuring machine (CMM) during dissection. A model was initially constructed in 3 dimensions, then reduced to a planar equivalent. Hill model parameters were estimated from architectural measurements. To further simplify calculations, muscles were lumped into 9 functional group equivalents based on mechanical actions.

#### Recording of passive limb kinematics

The right forelimb of four specimens was skinned and disarticulated at the scapula before establishment of *rigor mortis*. Self-tapping threaded rods were inserted into the scapula, humerus, ulna, and radius, and a 2-mm rod was driven through the medial-lateral axis of the paw. A plastic tubing elbow was fixed to each rod, and 4-mm reflective spheres mounted at the ends of the elbows and the end of each rod to produce bone-fixed marker triads. The limb was then manipulated within the field of view of a 6-camera Vicon motion capture system (Vicon Motion Systems Ltd, UK), with kinematic data collected at 100 Hz. Special attention was paid to moving each joint through a comfortable range of motion without imposing large forces or moments. Multiple recordings were made of each specimen, with some recordings emphasizing a single articulation and some making an accordion-like motion to change all joint angles simultaneously.

#### Measurement of moment arms

Eight forelimbs from four specimens were used for moment arm measurements. These limbs were skinned and disarticulated at the scapula following resolution of rigor mortis. For wrist moment arms, self-tapping, threaded bone pins driven through the radius and ulna were used to immobilize the specimen. Silk sutures were tied to the distal tendons of the prime wrist movers (flexor carpi ulnaris, FCU; extensor carpi ulnaris, ECU; flexor carpi radialis, FCR; and extensor carpi radialis, ECR) routed along the respective muscle bellies, wrapped around the shafts of rotary potentiometers, and fixed to 500-g suspended weights. A dual axis electrogoniometer was fixed to the paw and the forearm to record joint angles. The paw was then manipulated through flexion-extension and radial-ulnar deviation movements, while joint angles and tendon displacement were digitized. For elbow moment arms, the humerus was immobilized by bone pins, and sutures were connected to the triceps and biceps brachii tendons. Tendon-displacement vs joint angle curves were fitted to cubic polynomials and differentiated to give quadratic moment arms (An et al., 1983; Young et al., 1993).

#### Joint centers

To capture 3-D kinematics of the forelimb, we defined 7 rotational degrees of freedom or axes: 3 shoulder axes, elbow flexion-extension, radio-ulnar joint to capture pronation-supination, wrist dorsiflexion/plantarflexion, and wrist radial-ulnar deviation. Joint axes were identified by transforming the marker motion of each distal segment into a coordinate system defined by the marker triad of the adjacent proximal segment. An unconstrained optimization was used to determine the location (and axis orientation for hinge joints) of the joint center which minimized deviation of the transformed data from the surface (cylinder or sphere) defined by the joint. These joint centers were transferred into the specimen-specific mathematical models, and the recorded motions were played back for validation.

Because of substantial noise in the kinematic data, specific inclusion criteria were applied to the kinematic data sets. The spatial distribution of each point was required to exceed 20° to ensure that the joint was in fact moved during the specific manipulation. Furthermore, in some specimens, it was apparent that a set of marker triads moved relative to the bones at some point between collection of the kinematic data and digitization of individual points of interest (marker triads, bony landmarks, and muscle attachments), which prevented transformation of the joints proximal and distal to the corrupted marker into the anatomical coordinate system. This prevented accurate determination of joint centers for 2 elbow, 2 radio-ulnar, and 2 wrist joints. Thus, the preliminary axes for each joint were ultimately based on only two specimens, but two different specimens depending on the joint.

Preliminary joint definitions were further refined based on measured moment arms. Constrained optimization was used to minimize the sum square difference between the model and experimental moment arms for the biceps brachii and triceps muscles, subject to the constraints that the location of the elbow joint was not allowed to vary by more than 5 mm from the joint center as defined by the kinematic data and the orientation was not allowed to vary by more than 5°.

The radial-ulnar articulation was refined to reduce the variation in wrist moment arms with supination. We performed constrained optimization minimizing the difference between flexion-extension moment arms at 0° and 90° of supination while constraining the orientation of the radial-ulnar hinge axis that was allowed to vary by no more than 4° from the orientation obtained with kinematic data.

The wrist joint was refined by constrained optimization minimizing the difference between modeled and measured ECU, FCU, ECR and FCR moment arms where the location of the wrist joint was allowed to vary by no more than 5 mm from the joint center as defined by the kinematic data and the orientation was allowed to vary by no more than 10°.

#### Muscle attachments

Anatomical data were collected from six specimens. A set of rigid clamps was used to interlock the bone-mounted rods with the forelimb in a stance-like posture. The assembly was then immersed in 4% buffered formalin for 2-3 weeks. Care was taken not to displace or distort the marker triads, when present, which were used to align the kinematic data with the anatomical data. Following fixation, specimens were transferred to phosphate buffered saline (PBS) and stored at 4°C between dissection sessions.

To collect muscle attachment points, each specimen was mounted within the workspace of a 5-axis CMM (Microscribe, Immersion Technologies, USA). The locations of fixed points on the immobilization framework were recorded, followed by locations of kinematic triads, bony landmarks, and muscle origin, insertion, and via points. Each of these points was digitized at least three times, with the specimen removed from its mounting and moved to a different region of the CMM workspace between measurements. Superficial muscles obscured the connections of deeper muscles. As each superficial muscle was fully digitized, it was carefully dissected and reserved for architectural measurements. Each specimen required 4-6 layers of dissection, depending on posture.

The repeated digital point recordings were aligned to a common reference frame by equiform transformation (Veldpaus et al., 1988). This transformation relies on the fixed points in the immobilization framework, which were common to all the recordings of a particular specimen. The aligned anatomical points were then averaged to define a specimen-specific model. Because the specimens were fixed in different postures, the specimen-specific models were segmented, and the segments were aligned by equiform transformation. The aligned segments were then averaged to produce the final anatomical data set. Bony landmarks in each segment were used to define segmental coordinate systems.

#### Architectural measurements of muscles

Muscle-tendon units (MTU) from six specimens were used for architectural measurements. Each muscle was cleaned of excess connective tissue and fat and weighed. The length of MTU (*L^MTU^*) along its longest axis was measured with dial calipers and the surface pennation angle (*α*^*p*^), the angle between the apparent muscle line of action and predominant fiber direction, was measured with a goniometer. MTUs were then digested for 2-5 days in 10% nitric acid and rinsed several times in PBS. Bundles of muscle fibers were teased from the digested muscle, being careful to keep fascicles intact from proximal to distal tendons, and permanently mounted on glass slides. Between 3 and 5 fascicles were collected from different regions of each muscle. The length of each fascicle was measured with dial calipers. Sarcomere length of each bundle was determined by laser diffraction. The length of each fascicle was normalized to the optimal sarcomere length of 2.43 μm (Burkholder and Lieber, 2001), and the normalized fascicle lengths were averaged to give the optimal muscle fascicle length (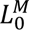). Physiological cross-sectional area (*PCSA*) was determined by the equation:

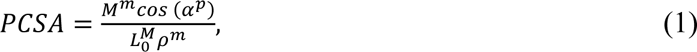

where *M*^*m*^ is the muscle mass, *α*^*p*^ is the pennation angle, 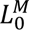 is the optimal fascicle length and *ρ*^*m*^ = 1.056 *g*/*cm*^3^ is muscle density (Mendez and Keys, 1960).

#### Muscle force model

The tendon force (*F*^*T*^) or the MTU force (*F*^*MTU*^) was computed based on the phenomenological Hill-type muscle model as a function of muscle activation (*A*, 0 ≤ *A* ≤ 1), muscle fascicle length (*L*^*M*^) and velocity (*V*^*M*^), pennation angle (*α*^*p*^) and passive series and parallel elastic elements (eq. 2-6) (He et al., 1991; Prilutsky et al., 2016). We simplified this model assuming that the series elastic component is rigid. The tendon slack length of each muscle was selected such that the maximum muscle fascicle length during the walking cycle was 105% of the optimal fiber length (Burkholder and Lieber, 2001):

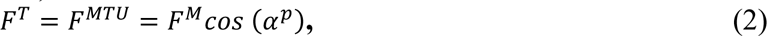

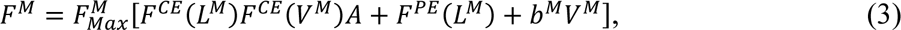

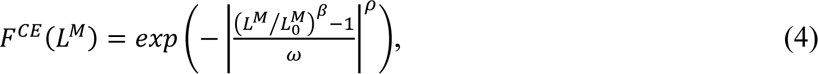

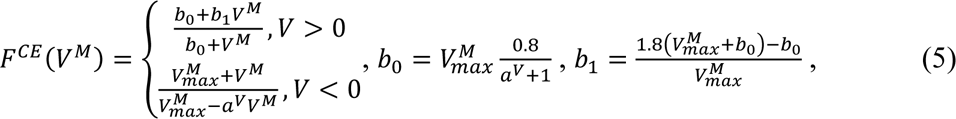

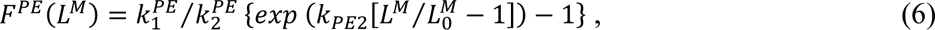

where *F*^*M*^ is muscle fascicle force; 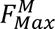 = 2.25 · *PCSA* is the maximum isometric muscle fascicle force at the optimum muscle fascicle length, *PCSA* is the muscle physiological cross-sectional area [in cm^2^] and constant 2.25 [in N/cm^2^] is muscle specific stress (Spector et al., 1980); *F*^*CE*^(*L*^*M*^) is the normalized isometric force-length relationship of the contractile element (muscle fascicles), eq. 4; *F*^*CE*^(*V*^*M*^) is the normalized force-velocity relationship of the contractile element, eq. 5; 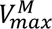 is the muscle maximum shortening velocity [in m/s]; *F*^*PE*^(*L*^*M*^) is the normalized force-length relationship of the muscle parallel elastic element, eq. 6. Parameters of eqs. 2 – 6 are summarized in Tables 5 and 6.

**Table 5.** Architectural properties of forelimb muscles.

| Muscle and primary joint action | Secondary joint action | Abbreviation | Muscle Mass, g | Optimal MTU length, mm | Optimal Fascicle Length, mm | Pennation Angle, deg | PCSA, cm<sup>2</sup> | Fiber type: SO, FOG, FG, % | $nV_{max}^M / s$ | Joint Radius R1, mm | Joint Radius R2, mm |
| --- | --- | --- | --- | --- | --- | --- | --- | --- | --- | --- | --- |
| **Shoulder protractors** |  | **Sh Prot** | **49.8** | **59.8** | **15.9** | **15** | **25.87** | **41.2,32.2,26.5** | **11.0** | **7.94** | — |
| Acromiodeltoideus | Shoulder abductor | AD | 2.79±0.47 | 59.1±15.2 | 13±2.9 | 31±14 | 2.04±0.6 | 21.9,78.1,0<sup>g</sup> | 11.0 | 2.0 | — |
| Coracobrachialis | Shoulder adductor | CB | 0.31±0.04 | 31.0±12.0 | 11±7.2 | 0±0 | 0.34±0.25 | 56,19,25<sup>a</sup> | 10.2 | 5.0 | — |
| Infraspinatus | Shoulder abductor | IF | 13.2±1.7 | 78.9±4.9 | 18.4±3.1 | 15±10 | 6.62±1.07 | 37,24,39<sup>a</sup> | 11.6 | 3.0 | — |
| Supraspinatus | - | SPS | 17.2±1.8 | 86.1±4.2 | 27.2±5.6 | 6±3 | 5.94±1.33 | 49,23,28<sup>a</sup> | 10.6 | 21.9 | — |
| Subscapularis | Shoulder adductor | SSC | 15.4±1.8 | 78.3±5.1 | 13.7±1.5 | 26±6 | 10.25±1.2 | 32,26,42<sup>a</sup> | 12.0 | 6.2 | — |
| Teres Minor | - | TN | 0.93±0.22 | 32.2±4.9 | 12.5±3.8 | 12±8 | 0.7±0.16 | 51,23,25<sup>a</sup> | 10.6 | 1.6 | — |
| **Shoulder retractors** |  | **Sh Ret** | **12.01** | **71.4** | **39.5** | **4** | **2.7** | **21.9,78.1,0** | **11** | **1.84** | — |
| Spinodeltoideus | Shoulder abductor | SD | 3.54±1.01 | 72.1±4.1 | 32.9±3.5 | 1±1 | 1.01±0.4 | 21.9,78.1,0<sup>g</sup> | 11 | 1.8 | — |
| Teres Major | - | TJ | 8.5±1.4 | 84.7±2.6 | 46.1±5.7 | 7±5 | 1.69±0.38 | 21.9,78.1,0<sup>b</sup> | 11 | 3.5 | — |
| **Shoulder Protr-Elbow Flex** |  | **Sh Prot-El Fl** | **5.9** | **70.5** | **25.1** | **10** | **2.27** | **20,21,59** | **11.9** | **10.0** | **5.30** |
| Biceps Brachii | - | BB | 5.9±1.0 | 81.0±8.7 | 25.1±7.6 | 10±10 | 2.27±0.81 | 20,21,59<sup>c</sup> | 11.9 | 10.0 | 5.3 |
| **Shoulder Retr-Elbow Ext** |  | **Sh Ret-El Ex** | **22.9** | **98.8** | **23.9** | **13** | **9.2** | **26,17,57** | **11.4** | **4.40** | **11.90** |
| Triceps Brachii Longus | - | TLONG | 22.9±3.9 | 93.1±8.1 | 23.9±7.1 | 13±11 | 9.2±2.75 | 26,17,57<sup>c</sup> | 11.4 | 4.4 | 11.9 |
| **Elbow extensors** |  | **El Ext** | **18.5** | **69.7** | **29.1** | **10** | **5.58** | **55.3,13.3,31.5** | **8.7** | **2.10** | — |
| Anconeus | Forearm pronator | ANC | 2.04±0.13 | 56.6±8.8 | 20.1±2.5 | 19±5 | 0.94±0.13 | 100,0,0<sup>c</sup> | 4.7 | 2.4 | — |
| Epitrochlearis | - | EPT | 1.95±0.51 | 66.5±8.5 | 30.1±3.6 | 0±0 | 0.59±0.16 | 26,17,57<sup>g</sup> | 11.4 | 2.1 | — |
| Triceps Brachii Lateralis | - | TLAT | 9.0±2.2 | 78.2±5.6 | 36.8±5.1 | 8±3 | 2.3±0.84 | 15,22,63<sup>c</sup> | 12.3 | 5.8 | — |
| Triceps Brachii Medialis | - | TMED | 5.6±0.4 | 83.8±7.9 | 29.2±3.8 | 11±4 | 1.75±0.21 | 80,14,6<sup>c</sup> | 6.4 | 8.4 | — |
| **Elbow flexors** |  | **El Fls** | **5.9** | **69.7** | **44.5** | **10** | **2.5** | **16.5,40.2,43.2** | **12.2** | **3.20** | — |
| Brachioradialis | Forearm supinator | BCD | 0.80±0.21 | 136.7±12.5 | 89.6±9.1 | 0±0 | 0.08±0.03 | 13,19,67<sup>a</sup> | 13.1 | 5.8 | — |
| Brachialis | - | BR | 3.47±0.35 | 88.6±6.0 | 32.4±4.4 | 10±4 | 0.99±0.22 | 25,24,51<sup>c</sup> | 11.4 | 6 | — |
| Pronator Teres | Forearm pronator | PT | 1.62±0.28 | 51.4±12.3 | 11.5±3.5 | 22±4 | 1.43±0.73 | 11.5,77.5,11.5<sup>b</sup> | 12 | 4.2 | — |
| **Elbow flexor-Wrist dorsiflexors** |  | **El Fl-Wr dFl** | **11.8** | **76.5** | **27.4** | **7** | **4.35** | **4,30.8,65.2** | **13.3** | **5.40** | **3.50** |
| Extensor Carpi Radialis | - | ECR | 3.88±0.66 | 98.3±19.4 | 47.3±35.3 | 15±0 | 0.97±0.59 | 4,41.9,53.9<sup>d</sup> | 13.1 | 5.5 | 3.5 |
| Extensor Digitorum Communis | - | EDC | 1.98±0.17 | 85.8±3.7 | 22.5±6.7 | 5±4 | 0.85±0.21 | 4,28,68<sup>e</sup> | 13.3 | 5.3 | 6.1 |
| **Wrist plantarflexors** |  | **Wr pFl** | **65.4** | **79.8** | **13.2** | **8** | **36.97** | **20.1,30.4,49.4** | **11.7** | **2.70** | — |
| Flexor Carpi Radialis | - | FCR | 1.26±0.17 | 83.4±15.7 | 14.3±2.3 | 6±1 | 0.85±0.14 | 38.1,29.4,32.3<sup>f</sup> | 10.1 | 2.7 | — |
| Flexor Carpi Ulnaris | - | FCU | 4.01±0.32 | 104.0±2.6 | 11.1±1.1 | 11±2 | 3.3±0.31 | 42.4,24.2,33.4<sup>f</sup> | 9.7 | 5.6 | — |
| Flexor Digitorum Profundus | - | FDP | 9.27±2.00 | 103.9±6.1 | 21.5±2.7 | 7±12 | 3.97±1.07 | 8.1,34.6,57.2<sup>f</sup> | 12.8 | 5.5 | — |
| Flexor Digitorum Superficialis | - | FDS | 0.18±0.05 | 29.1±4.3 | 5.85±2.2 | 3±3 | 0.29±0.07 | 26.8,29.2,43.9<sup>g</sup> | 11.1 | 2.7 | — |
| Palmaris Longus | - | PL | 2.61±0.55 | 101.8±22.2 | 10.9±1.4 | 14±6 | 2.36±0.61 | 18.7,28.7,52.6<sup>f</sup> | 11.9 | 6.3 | — |
| **Wrist dorsiflexors** |  | **Wr dFl** | **1.11** | **78.4** | **21** | **1** | **0.25** | **2,55,43** | **13.2** | **2.40** | — |
| Extensor Pollicis Longus | - | EPL | 0.55±0.07 | 86.1±8.1 | 21±6 | 1±1 | 0.25±0.08 | 2,55,43 | 13.2 | 2.4 | — |

**Table 6.** Selected parameters of the muscle model.

| Muscle force relationship | Equation | Parameter | Value |
| --- | --- | --- | --- |
| Force-velocity relationship of passive muscle | 3 | $b^M$ | $0.02 \text{ N} \cdot \text{s}/\text{m}$ |
| Normalized isometric force-length relationship of contractile component, $F^{CE}(L^M)$ | 4 | $\omega$<br>$\rho \text{ (for } L^T/L_0^M \leq 1)$<br>$\rho \text{ (for } L^T/L_0^M > 1)$<br>$\beta$ | 0.55<br>6.0<br>3.0<br>1.55 |
| Normalized force-length relationship of passive elastic element, $F^{PE}(L^M)$ | 6 | $k_1^{PE}$<br>$k_2^{PE}$ | 0.0075<br>11.6 |

#### Afferent activity model

Firing rates of muscle spindle and Golgi tendon organ afferents during walking were estimated from the computed muscle fascicle length and velocity as well as MTU force using the regression equations developed in (Prochazka and Gorassini, 1998a; Prochazka, 1999). The firing rates of spindle group Ia and II and Golgi tendon organ group Ib afferents were computed as follows:

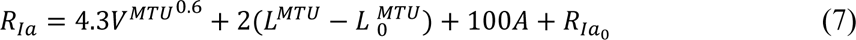

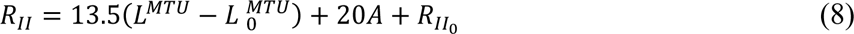

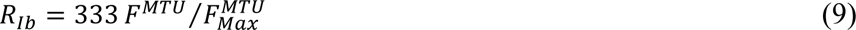

where *R*_*Ia*_ is the firing rate of muscle spindle group Ia afferents (in *HZ*); *V*^*MTU*^ is MTU velocity (in mm/s); 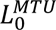 is optimal MTU length (in mm); *R*_*Ia*_0__ is the mean of *R*_*Ia*_ over the cycle (in Hz); *R*_*II*_ is the firing rate of muscle spindle group II afferents (in Hz); *R*_*II*_0__ is the mean of *R*_*II*_ over the cycle (in Hz); *R*_*Ib*_ is the firing rate of Golgi tendon organ group Ib afferents (in Hz); 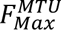 = 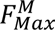 *cos* (*α*^*p*^) is the maximum isometric MTU force (see eq. 2).

#### Reduction to planar model

We fitted a plane into 3D coordinates of attachment and via points of all studied forelimb muscles (Figure 1A, 3D view; Table S1) by computing the first two principal components of the 3D coordinate data. We defined this plane as the forelimb sagittal plane. We projected muscle attachment and via points on this plane. We also computed the intersection points of the joint axes with the sagittal plane and defined these intersection points as joint centers in this plane. The sagittal plane representation of the forelimb segments (connecting joint centers) and muscles (connecting attachment and via points) is shown in Figure 1A, Sagittal view (Table S2). A minimum moment arm, or wrapping circle, was defined for each joint to prevent muscles from crossing the joint center. All further analysis is based on this 5 degrees-of-freedom (DOF) model, in which the first two DOFs represent the horizontal and vertical coordinates of the shoulder marker (joint), and the other three DOFs represent the angles at the shoulder, elbow, and wrist joints, as defined in Fig. 1, Sagittal view.

**Figure 1.**
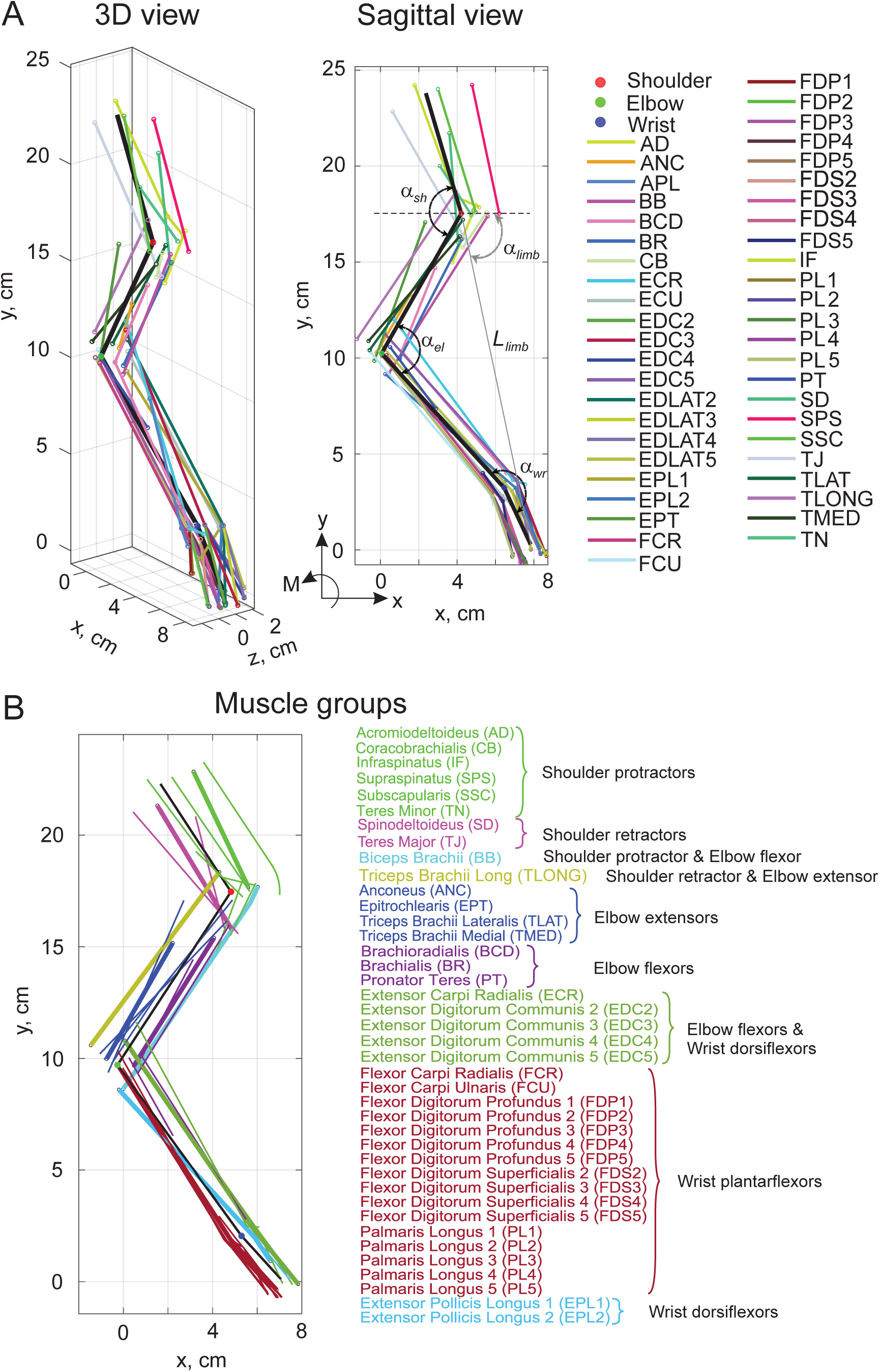
Schematic of cat forelimb musculoskeletal model. Black lines designate the scapular, upper arm, forearm, and carpals. Filled color circles indicate locations of the shoulder, elbow, and wrist centers. α_sh_, α_el_, and α_wr_ define shoulder, elbow, and wrist joint angles. Open small circles indicate origin, via points, and attachment of muscle-tendon units (MTUs). Color lines connecting the origin and attachment points correspond to forelimb MTUs. **A**: 3D and 2D sagittal plane views of cat forelimb with 46 MTUs. Thin gray line connecting the shoulder center and the most distal carpals point define the forelimb length (*L_limb_*); angle α*_limb_* defines limb orientation. **B**: Results of the cluster analysis of mechanical actions of 40 MTUs based on their maximum moment of force at joints in the sagittal plane. Six MTUs whose major mechanical action is outside the sagittal plane are excluded from cluster analysis. Thin and thick lines of the same color correspond to individual MTUs and their equivalent MTU with the same mechanical action at a joint, respectively (see text for details).

#### Calculations of MTU lengths and moment arms during locomotion

We computed the instantaneous length of each MTU in locomoting cats from the experimental joint angles and morphological parameters of the MTU and joints. The morphological parameters included the distance of the MTU attachment point on the proximal (α_1_) and distal (α_2_) segment to the joint center, the angle formed by the line from the attachment point to the joint center and the proximal (*φ*_1_) and distal (*φ*_2_) segments, the radius of the joint spanned by a one-joint muscle (*R*_1_) or the radii of the proximal (*R*_1_) and distal (*R*_2_) joints spanned by a two-joint muscle (Fig. 2). Additional geometric parameters necessary for calculating the MTU length and the corresponding equations are shown in Fig. 2. The equation parameters depend on the location of the MTU path with respect to one or two joints spanned by the muscle and on whether the MTU path wraps around the joints. Additional information about calculations of MTU lengths considering wrapping around joints and geometric parameters can be found in Table 5 and Supplement 3. The moment arm of the *i*-th MTU with respect to the *j*-th joint, *r_i,j_*, was computed as the derivative of the MTU length over the joint angle.

**Figure 2.**
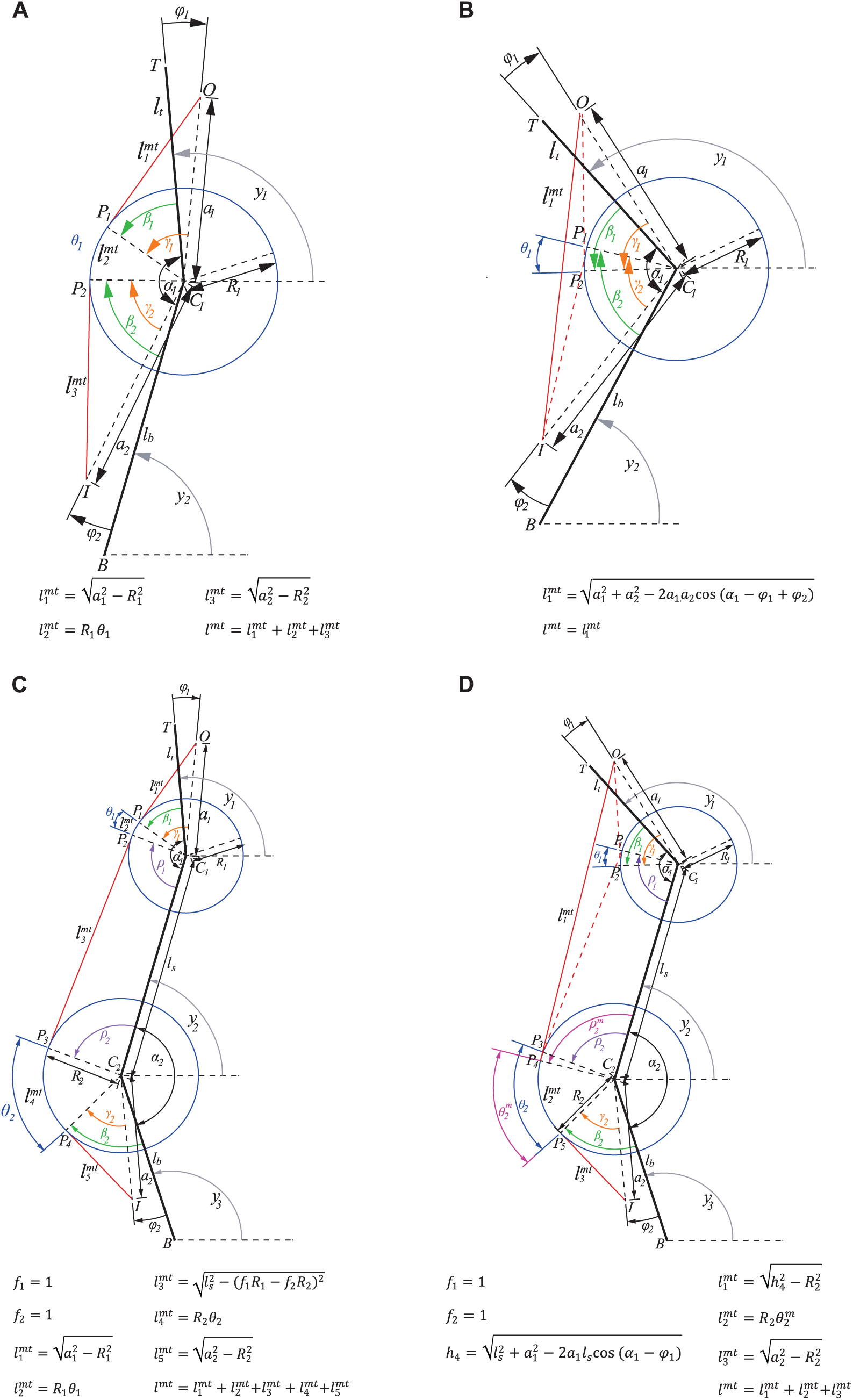
Schematics demonstrating calculations of the path, length, and moment arms of one- and two-joint MTUs as functions of the joint angles and geometric parameters of the MTU and joint. O and I are points of MTU origin and insertion; α_1_ and α_2_ are distances from the joint center to O and I, respectively; φ_1_ and φ_2_ are angles formed by the limb segment and lines from the joint center to O and I, respectively; R_1_ and R_2_ are joint radii. If the MTU path is not in contact with joint surface, defined by the circle around the joint center, MTU length is defined as the distance between O and I. If the MTU path wraps around the joint surface, the total MTU length is the sum of the three lengths: 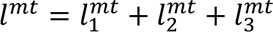 (see figure for definitions of these and other parameters). MTU moment arm at a joint is computed as the shortest distance from the MTU path to the joint center. **A**, **B**: Path definition for one-joint MTUs located posterior to the joint center with wrapping or without wrapping around the joint. **C**, **D**: Path definition for two-joint MTUs located posterior to the joint centers with different combinations of wrapping around the two joints.

#### Reduction to functional groups

We classified muscles based on their maximum moment of force produced in the walking cycle. The maximum moment of force was computed for each percent of the cycle as the product of the MTU moment arm and MTU force assuming maximum muscle activation (*A* = 1). MTUs were classified into functional groups using the K-means clustering algorithm of MATLAB (MathWorks, Inc., USA). The equivalent mass and PCSA of each functional group were computed as the sum of masses and PCSAs of the MTUs within each cluster. The other morphological parameters (Fig. 2, Table 7) of each functional group were determined by minimizing the difference between the total maximum moment of MTUs in a cluster and the moment of the equivalent MTU produced during the walking cycle (optimization function *fmincon* in MATLAB):

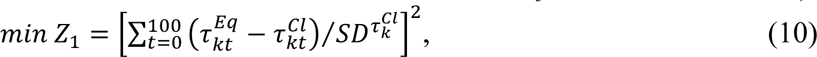

**Table 7.** Geometric parameters of 9 forelimb equivalent MTUs representing sagittal plane actions of the corresponding individual MTUs with the same actions at the joints.

| Muscle | $a_1$, cm | $a_2$, cm | $\phi_1$, rad | $\phi_2$, rad | $R_1$, cm | $R_2$, cm |
| --- | --- | --- | --- | --- | --- | --- |
| Shoulder protractor | 5.64 | 0.795 | -0.277 | 2.392 | 0.794 | - |
| Shoulder retractor | 5.08 | 1.67 | 0.128 | 0.595 | 0.184 | - |
| Shoulder protractor –<br>Elbow flexor | 1.22 | 1.11 | -1.969 | -0.400 | 1.00 | 0.53 |
| Shoulder retractor –<br>Elbow extensor | 1.02 | 1.49 | -0.046 | -2.832 | 0.44 | 1.19 |
| Elbow extensor | 6.00 | 0.56 | 0.155 | -2.691 | 0.21 | - |
| Elbow flexor | 7.15 | 0.74 | -0.070 | 0.899 | 0.32 | - |
| Elbow flexor –<br>Wrist dorsiflexor (OV) | 1.14 | 9.51 | 0.287 | 0.065 | 0.54 | 0.35 |
| Elbow flexor –<br>Wrist dorsiflexor (VI) | 0.61 | 3.33 | -1.531 | 0.126 |  |  |
| Wrist plantarflexor (OV) | 0.64 | 9.63 | -0.047 | -0.052 | 0.27 | - |
| Wrist plantarflexor (VI) | 0.52 | 2.89 | 1.282 | -0.163 |  |  |
| Wrist dorsiflexor (OV) | 1.11 | 9.89 | -0.571 | 0.044 | 0.24 | - |
| Wrist dorsiflexor (VI) | 0.601 | 1.71 | -2.333 | 0.115 |  |  |

Note. The geometric parameters are explained in Fig. 2. The MTU paths of the Elbow flexor – Wrist dorsiflexor, Wrist plantarflexor, and Wrist dorsiflexor consist of two lines connected by a via point (designated by V). The first line connects the point of origin (O) with V, and the second line connects V with the point of insertion (I).

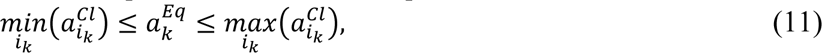

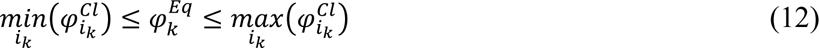

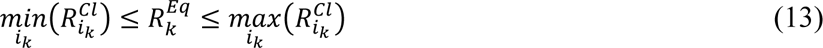

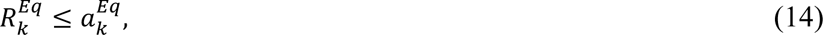

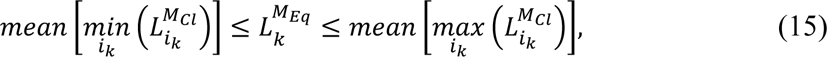

where 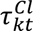 and 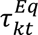 are the total moment of all MTUs in *k*-th cluster at the *t*-th instant of the normalized cycle time and the moment of the equivalent MTU substituting the total moment of the same MTU cluster at time instant *t*, respectively; 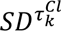 is the standard deviation of the total MTU cluster moment computed across all MTUs in the cluster at time *t*; 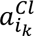 is the distance from an attachment point to the joint center of *i*-th MTU of cluster *k* (note that each MTU has two attachment points, Fig. 2); 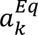 is the distance from an attachment point to the joint center of the equivalent MTU for cluster *k*; 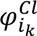 is the angle between the segment line and the line between the joint center and an attachment point of *i*-th MTU of cluster *k* (each MTU has two such angles, Fig. 2); 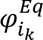 is the angle between the segment line and the line between the joint center and an attachment point of the equivalent MTU of cluster *k*; 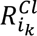 is the joint radius of individual *i*-th MTU in *k*-th cluster at a joint this MTU spans; 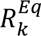 is the joint radius of the equivalent MTU for cluster *k* at a joint spanned by this MTU; 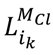 and 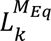 are length of *i*-th MTU in cluster *k* and length of the equivalent MTU for cluster *k* in the walking cycle.

### Gait analysis and inverse dynamics

#### Strategy

Overground walking kinematics were recorded by infrared motion capture and ground reaction forces measured by force platforms. Muscle activations during walking were estimated by static optimization with a minimum fatigue cost function (Crowninshield and Brand, 1981). Estimation was performed for both the fully redundant, 40-MTU model and the simplified, 9-functional group model.

#### Recording and analysis of forelimb locomotor kinematics and ground reaction forces

Seven cats (Tables 2 and 3) were trained to walk at self-selected speeds on a Plexiglas enclosed walkway with three embedded small force platforms (Bertec Corporation, USA) using food rewards and affection. After training for 2-4 weeks, animals were sedated (dexmedetomidine, 40–60 μg/kg, i.m.) and shaved. In the subsequent 2 weeks, we recorded full-body kinematics – 3D coordinates of 28 small reflective markers on major hindlimb and forelimb joints, the scapulas, the pelvis and head (6-camera Vicon motion capture system, sampling frequency 120 Hz). We also recorded the three components of ground reaction forces applied to each hindpaw and forepaw as well as the coordinates of the force vector application at a sampling frequency of 360 Hz (Farrell et al., 2014; Klishko et al., 2014). We attached the reflective markers using double sided adhesive tape on the following bone landmarks of the left and right forelimbs: intersection of the vertebral border and tuberosity of the spine on the scapula (the point approximating halfway between scapula cranial and caudal angles), humerus greater tubercle, humerus lateral epicondyle, ulna styloid process, lateral metacarpophalangeal joint (MCP) and distal phalanx of the 5^th^ digit. After low-pass filtering recorded marker coordinates (4^th^ order, zero-lag Butterworth filter, cutoff frequency 5-7 Hz), the coordinates of the elbow joint were recalculated by triangulation using coordinates of the humerus greater tubercle and ulna styloid process and segment lengths of the upper arm and forearm (Table 2) to reduce skin movement artefacts.

We computed the moment of force (resultant muscle moment, *τ*_*j*_) at *j*-th joint in the sagittal plane using 2D Newton-Euler equations of motion of each forelimb segment distal to the joint, the recorded marker coordinates and ground reaction forces as well as the segment inertia parameters (mass, center of mass location, moment of inertia with respect to the center of mass, Table 3 (Prilutsky et al., 2005; Farrell et al., 2014)).

We identified individual locomotor cycles from successive swing onsets. The swing onset (stance offset) and swing offset (stance onset) were defined based on the relative horizontal displacement of the forepaw with respect to the shoulder, the method demonstrating the smallest random error of phase detection for the hindlimb (Pantall et al., 2012).

#### Implantation of EMG electrodes and recording and analysis of EMG activity

EMG activity of selected forelimb muscles was recorded in 16 cats walking on a tied-belt treadmill with speed of 0.4 m/s. These animals were also used in other studies addressing different scientific questions (Lecomte et al., 2022; Merlet et al., 2022; Audet et al., 2023; Lecomte et al., 2023; Mari et al., 2023; Harnie et al., 2024; Mari et al., 2024). Implantations of EMG electrodes and recording procedures were described in those studies, therefore only a brief account is provided here. Before surgery, the cat was sedated with an intramuscular injection of a cocktail containing butorphanol (0.4 mg/kg), acepromazine (0.1 mg/kg) and glycopyrrolate (0.01 mg/kg) and inducted with another intramuscular injection (0.05 ml/kg) of ketamine (2.0 mg/kg) and diazepam (0.25 mg/kg) in a 1:1 ratio. The fur overlying the forelimbs and top of the head was shaved and the skin cleaned with chlorhexidine soap. Implantation surgeries were performed under aseptic conditions and general anesthesia with isoflurane (1.5-3.0%) and O_2_ delivered with a flexible endotracheal tube. The depth of anesthesia was confirmed by testing a withdrawal response to applying pressure to a paw. Body temperature was monitored and maintained at 37 ± 0.5°C using a water-filled heating pad under the animal and an infrared lamp placed ∼50 cm above it. Pairs of Teflon-insulated multistrain fine wires (AS633; Cooner Wire Co., Chatsworth, CA, USA) were subcutaneously passed from two head-mounted 34-pin connectors (Omnetics Connector Corp., Minneapolis, MN, USA) to the muscles of interests. Two wires, stripped of 1–2 mm of insulation, were sewn into the belly of selected forelimb muscles for bipolar recordings. The head-mounted connectors were fixed to the skull using dental acrylic and four to six screws. Seven forelimb muscles were implanted bilaterally: supraspinatus (shoulder protractor), biceps brachii (shoulder protractor and shoulder flexor), triceps brachii long head (shoulder retractor and elbow extensor), triceps brachii lateral head (elbow extensor), pronator teres (shoulder flexor and forearm pronator), brachialis (shoulder flexor), and extensor carpi radialis (elbow flexor and wrist dorsiflexor). Electrode placement was verified during surgery by stimulating each muscle through the appropriate head connector channel. The skin was closed using subcuticular sutures (monocryl 4–0, Ethicon) followed by cutaneous sutures (monocryl 3–0, Ethicon). At the end of surgery, an antibiotic (cefovecin, 8 mg/kg) and a fast-acting analgesic (buprenorphine, 0.01 mg/kg) were injected subcutaneously. A fentanyl (25 µg/h) patch was taped to the back of the animal 2-3 cm rostral to the base of the tail for prolonged analgesia, which was removed 4-5 days later. After surgery, cats were placed in an incubator and closely monitored until they regained consciousness. Another dose of buprenorphine was administered ∼7 hours after surgery. At the end of experiments, cats were anaesthetized with isoflurane (1.5–3.0%) and O_2_ before receiving a lethal dose of pentobarbital (120 mg/kg) through the left or right cephalic vein. Cardiac arrest was confirmed using a stethoscope to determine the death of the animal.

During experiments, EMG signals were pre-amplified (×10, custom-made system), bandpass filtered (30–1,000 Hz) and amplified (100–5,000×) using a 16-channel amplifier (model 3500; AM Systems, Sequim, WA, USA). EMG data were digitized (5,000 Hz) with a National Instruments (Austin, TX, USA) card (NI 6032E), acquired with custom-made acquisition software and stored on computer. Recorded EMG signals were full-wave rectified and low-pass filtered (Butterworth, zero lag 4^th^ order filter, 10 Hz cutoff frequency). Locomotion cycle durations were identified for each forelimb as the time between swing onset and stance offset events. These time events were obtained from bilateral video recordings made by two cameras (Basler AcA640-100 G) at 60 frames per second with a spatial resolution of 640 x 480 pixels. A custom-made program (LabView, National Instruments, USA) acquired the video images and synchronized them with EMG data. Low-pass filtered EMG signals were normalized to the peak filtered EMG across all cycles within the cat and muscle and then time-normalized to the duration of the cycle.

#### Estimation of muscle activation

To better understand forelimb locomotor functions, we computed patterns of muscle activation during the walking cycle using static optimization (Crowninshield and Brand, 1981; Prilutsky et al., 1997; Anderson and Pandy, 2001). Subject-specific models were generated for each animal by segment-length scaling of the planar model. The static optimization problem for each normalized cycle time instant was formulated as follows:

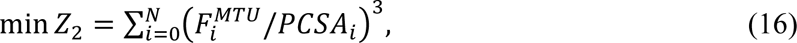

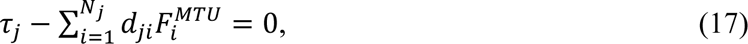

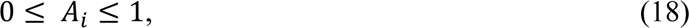

where *Z*_2_ is a cost function of minimum fatigue (Crowninshield and Brand, 1981); *j* indicates joint (*j*=1, 2, 3 are wrist, elbow, and shoulder, respectively); *τ*_*j*_ is the resultant moment of force averaged across all cycles of all cats for which we recorded kinematics and ground reaction forces as well as determined the segment inertial properties (Table 3); *d_ji_* is the mean moment arm of the *i*-th MTU with respect to the *j*-th joint determined from the model geometry, Fig. 2; 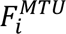 is the force of *i*-th MTU that depends on muscle activation *A_i_*, muscle fascicle pennation angle, length and velocity (eqs. 2-6). We used constrained optimization (function *fmincon* in MATLAB) to find the optimal muscle activations at each time instant of the normalized cycle time starting from random initial activation values. The optimal activations were found for the two forelimb models, with all 40 MTU units and with the reduced number of equivalent MTU units each representing the functional cluster.

#### Estimation of spatiotemporal activity of cervical motoneuronal pools and their proprioceptive inputs during walking

Muscle nerves originating in cervical segments of the spinal cord consisting of axons from motor and proprioceptive sensory neurons innervate forelimb MTUs. There is a well-established anatomical organization of motoneuronal pools of forelimb muscles along the rostral-caudal direction of cervical spinal segments with proprioceptive afferents connecting to motoneurons of the parent muscles and, to a lesser extent, their synergists (Sterling and Kuypers, 1967b, a; Fritz et al., 1986a; Fritz et al., 1986b; Levine et al., 2012). This anatomical organization allows for evaluation of spatiotemporal activity of motoneurons during a locomotor cycle based on recorded or computed muscle activity as was done for the lumbar motor pools of locomoting cats (Yakovenko et al., 2002), for the entire spinal cord of humans (Ivanenko et al., 2006) and for proprioceptive inputs to cervical spinal segment during arm reaching and grasping in monkeys (Kibleur et al., 2020).

We used this approach to map the computed activation of forelimb MTUs (eqs. 16-18) and their proprioceptive activity (eqs. 7-9) during locomotion on to cervical spinal segments. We used available data on the distribution of motor pools of cat forelimb muscles in spinal segments C3 through T1 from (Fritz et al., 1986a; Fritz et al., 1986b; Horner and Kummel, 1993). Because the cat data were missing for the BB, TLONG, FDS, and EPL MTUs (MTU abbreviations are defined in Fig. 1 and Table 5) in the mentioned studies, we approximated their motor pool localizations using data on arm motor pools in monkeys (Jenny and Inukai, 1983; Kibleur et al., 2020). Since we could not find reliable data on MTUs CB, ANC, EPT, TLAT, TMED, and BR, we approximated their motor pool localizations by the mean values of their synergistic MTUs. To quantify the distribution of motor pools along the spinal segments from published distribution histograms, we digitized them using GetData Graph Digitizer 2.26 software (Informer Technologies, Inc.) and then computed the total histogram area and the area corresponding to each spinal segment. The resulting proportions of motoneurons for each MTU in each cervical spinal segment are shown in Table 8. Assuming that motoneuronal pools of forelimb muscles and their proprioceptive inputs have the same rostro-caudal spinal distribution (Sterling and Kuypers, 1967b, a; Levine et al., 2012), we computed motor and sensory neuronal activities exiting and entering each spinal segment at each percent of the locomotor cycle as follows (Kibleur et al., 2020):

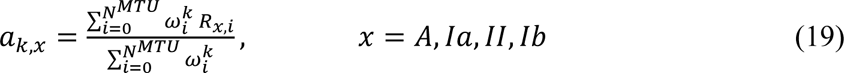

**Table 8.**
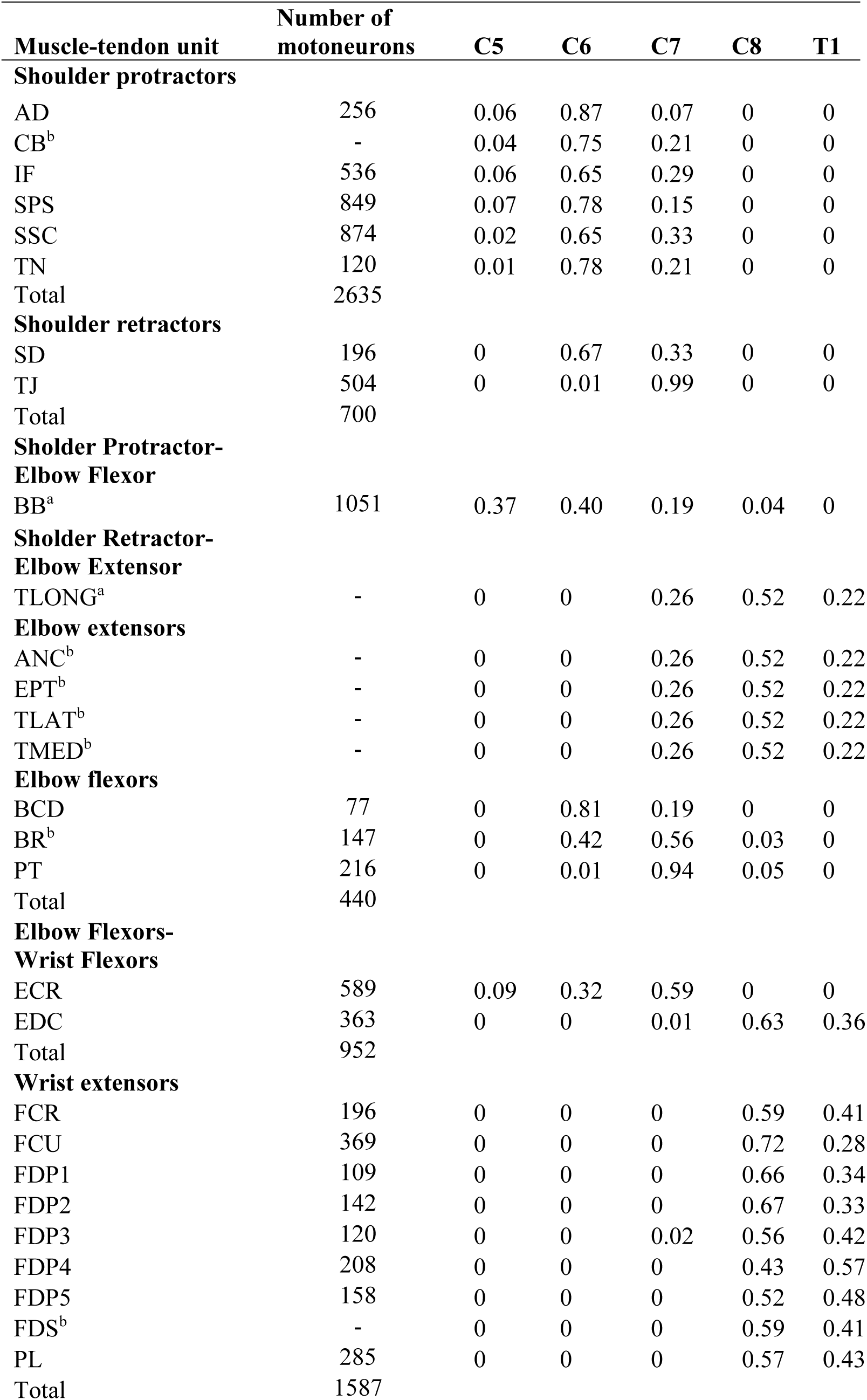
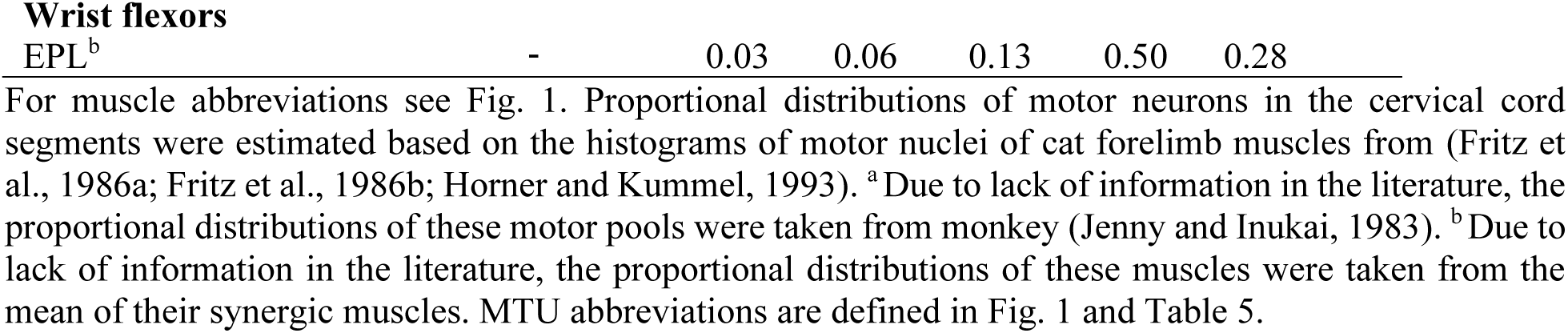
Proportional distribution of the motor neurons of forelimb muscles in spinal cord segments.

| **Muscle-tendon unit** | **Number of motoneurons** | **C5** | **C6** | **C7** | **C8** | **T1** |
| --- | --- | --- | --- | --- | --- | --- |
| **Shoulder protractors** |  |  |  |  |  |  |
| AD | 256 | 0.06 | 0.87 | 0.07 | 0 | 0 |
| CB<sup>b</sup> | - | 0.04 | 0.75 | 0.21 | 0 | 0 |
| IF | 536 | 0.06 | 0.65 | 0.29 | 0 | 0 |
| SPS | 849 | 0.07 | 0.78 | 0.15 | 0 | 0 |
| SSC | 874 | 0.02 | 0.65 | 0.33 | 0 | 0 |
| TN | 120 | 0.01 | 0.78 | 0.21 | 0 | 0 |
| Total | 2635 |  |  |  |  |  |
| **Shoulder retractors** |  |  |  |  |  |  |
| SD | 196 | 0 | 0.67 | 0.33 | 0 | 0 |
| TJ | 504 | 0 | 0.01 | 0.99 | 0 | 0 |
| Total | 700 |  |  |  |  |  |
| **Sholder Protractor-Elbow Flexor** |  |  |  |  |  |  |
| BB<sup>a</sup> | 1051 | 0.37 | 0.40 | 0.19 | 0.04 | 0 |
| **Sholder Retractor-Elbow Extensor** |  |  |  |  |  |  |
| TLONG<sup>a</sup> | - | 0 | 0 | 0.26 | 0.52 | 0.22 |
| **Elbow extensors** |  |  |  |  |  |  |
| ANC<sup>b</sup> | - | 0 | 0 | 0.26 | 0.52 | 0.22 |
| EPT<sup>b</sup> | - | 0 | 0 | 0.26 | 0.52 | 0.22 |
| TLAT<sup>b</sup> | - | 0 | 0 | 0.26 | 0.52 | 0.22 |
| TMED<sup>b</sup> | - | 0 | 0 | 0.26 | 0.52 | 0.22 |
| **Elbow flexors** |  |  |  |  |  |  |
| BCD | 77 | 0 | 0.81 | 0.19 | 0 | 0 |
| BR<sup>b</sup> | 147 | 0 | 0.42 | 0.56 | 0.03 | 0 |
| PT | 216 | 0 | 0.01 | 0.94 | 0.05 | 0 |
| Total | 440 |  |  |  |  |  |
| **Elbow Flexors-Wrist Flexors** |  |  |  |  |  |  |
| ECR | 589 | 0.09 | 0.32 | 0.59 | 0 | 0 |
| EDC | 363 | 0 | 0 | 0.01 | 0.63 | 0.36 |
| Total | 952 |  |  |  |  |  |
| **Wrist extensors** |  |  |  |  |  |  |
| FCR | 196 | 0 | 0 | 0 | 0.59 | 0.41 |
| FCU | 369 | 0 | 0 | 0 | 0.72 | 0.28 |
| FDP1 | 109 | 0 | 0 | 0 | 0.66 | 0.34 |
| FDP2 | 142 | 0 | 0 | 0 | 0.67 | 0.33 |
| FDP3 | 120 | 0 | 0 | 0.02 | 0.56 | 0.42 |
| FDP4 | 208 | 0 | 0 | 0 | 0.43 | 0.57 |
| FDP5 | 158 | 0 | 0 | 0 | 0.52 | 0.48 |
| FDS<sup>b</sup> | - | 0 | 0 | 0 | 0.59 | 0.41 |
| PL | 285 | 0 | 0 | 0 | 0.57 | 0.43 |
| Total | 1587 |  |  |  |  |  |
| **Wrist flexors** |  |  |  |  |  |  |
| EPL<sup>b</sup> | - | 0.03 | 0.06 | 0.13 | 0.50 | 0.28 |

where *a_k,x_* is activity of type *x* in the *k*-th spinal segment; *R_x,i_* is the normalized motoneuronal activity or the firing rate of the proprioceptive afferents of type *x* of the *i*-th MTU, and 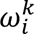 is the proportion of *i*-th MTU’s motoneurons in or its afferents projecting to the *k*-th spinal segment.

### Statistical analysis

Comparisons between the functional groups of MTUs were made for individual morphological characteristic using a one-way analysis of variance (Statview 4.0, Abacus concepts, Inc., Berkeley, CA), and in the case of a significant statistical value (p < 0.05), a Fisher’s protected least significant difference (PLSD) post-hoc test was performed between pairs of functional groups.

## RESULTS

### Forelimb morphology

The 3D musculoskeletal model of the cat forelimb and its 2D sagittal plane version are shown in Fig. 1A. The model includes four body segments (scapula, upper arm, forearm, and carpals), three joints (shoulder, elbow, and wrist), and MTUs of 46 muscles. For analysis of a muscle’s motor and sensory functions during locomotion in the sagittal plane, we selected 40 MTUs (Fig. 1B), because the remaining six MTUs have negligible moments of force production in the sagittal plane. The six muscles excluded from the sagittal plane analysis were abductor pollicis longus (APL, wrist abductor), extensor carpi ulnaris (ECU, wrist dorsiflexor), and four compartments of extensor digitorum lateralis (EDLAT2, EDLAT3, EDLAT4, EDLAT5, wrist dorsiflexors).

Table 5 shows forelimb architectural measures of muscles of the 2D model (Fig. 1B). The forelimb MTUs in Table 5 are classified by functional groups and vertically arranged from proximal to distal. Qualitatively, the forelimb is more complex than the hindlimb, with substantially more and denser connective tissue and compartmentalization. The cat forelimb possesses many complex, multi-pennate muscles (see Fig. 1A, Table 5), unlike the more fusiform, uni-pennate muscles typically found in the cat hindlimb (Sacks and Roy, 1982). In addition, forelimb MTUs tend to have shorter external tendons, with fibers inserting near the distal joint.

Figure 3 shows select morphological characteristics of forelimb MTUs combined with respect to their actions in 3D (Fig. 1A). Muscle masses ranged from 0.18 ± 0.05 g for FDS to 22.9 ± 3.9 g for TLONG. As shown in Fig. 3A, muscle mass varied significantly among functional groups (p < 0.0001), with greater and smaller masses concentrated in the proximal and distal limb, respectively. The shoulder protractors had the greatest mass (17.2 ± 1.8 g), followed by the shoulder adductors (10.3 ± 7.9 g) and elbow extensors (8.8 ± 8.5 g). These groups were significantly larger than the forearm supinator (0.86 ± 0.16 g), forearm pronator (1.86 ± 0.29 g), wrist dorsiflexor (1.70 ± 0.93 g) and wrist abductor (1.82±0.27 g) groups.

**Figure 3.**
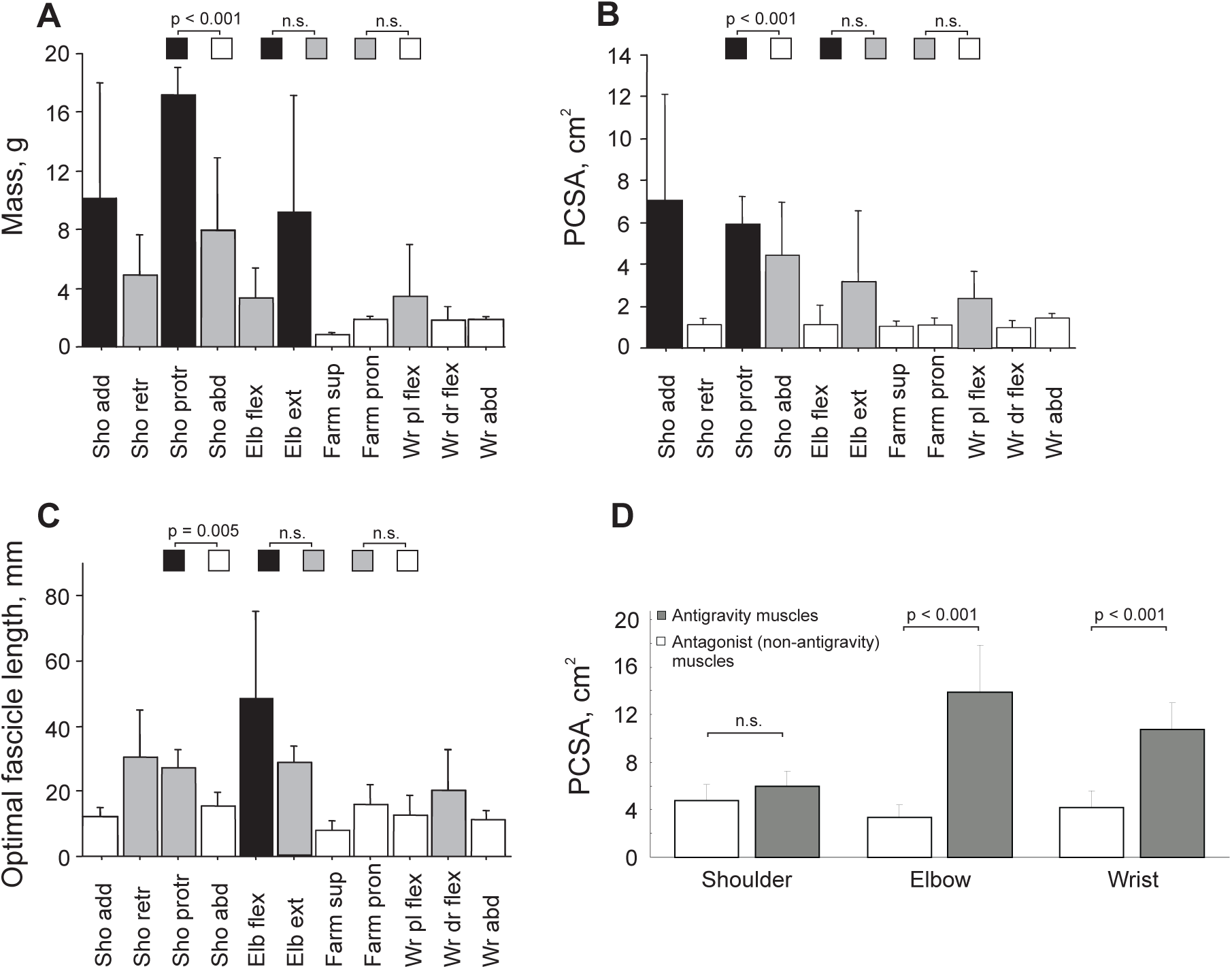
Morphological characteristics (mean ± SD) of 46 MTUs of the 3D forelimb model (see Fig. 1A). **A**, **B**, **C**: Mass, physiological cross-sectional area (PCSA), and optimal fascicle length, respectively, of MTU groups classified by their mechanical action in 3D. Values indicated by black and white bars are significantly different from each other (p ≤ 0.005). Gray bars indicate values that are statistically indistinguishable from black and white bars as well as from each other. Sho, Elb, Farm, Wr indicate shoulder, elbow, forearm, and wrist, respectively; add and abd, adductors and abductors; retr and protr, retractors and protractors; flex and ext, flexors and extensors; sup and pron, supinators and pronators; pl and dr, plantar and dorsi. **D**: Combined PCSA of antigravity muscles and their antagonists for the shoulder, elbow, and wrist (mean ± SD). P < 0.001 indicates significant difference, n.s. indicates no significant difference.

The calculated physiological cross-sectional area ranged from 0.08 ± 0.03 cm^2^ (BCD) to 10.25 ± 1.19 cm^2^ (SSC) (Table 5) with a proximo-distal gradient that was qualitatively more clearly defined than that of muscle mass (Fig. 3B). The shoulder adductors and protractors had the numerically largest PCSAs (6.94 ± 5.20 cm^2^ and 5.94 ± 1.33 cm^2^, respectively), roughly 3 times that of the majority of muscle groups, including shoulder retractors (1.13 ± 0.53 cm^2^), forearm supinators (1.09 ± 0.31 cm^2^) and pronators (1.15 ± 0.51 cm^2^), wrist abductors (1.43 ± 0.25 cm^2^), plantarflexors (2.21 ± 1.57 cm^2^), and dorsiflexors (0.90 ± 0.48 cm^2^). Antigravity muscles had higher PCSAs than their antagonists (Fig. 3D; p < 0.0001, one-way ANOVA). The ratio of combined anti-gravity to antagonist PCSA was 5:4 for shoulder muscles and approximately 4:1 at the elbow and 5:2 at the wrist (Fig. 3D). This supports the hypothesis that weight-bearing functional groups possess greater force-producing capacity than their antagonist.

The optimal fascicle length varied from 5.85 ± 2.19 mm (FDS) to 89.6 ± 9.1 mm (BCD) (Table 5), with the elbow flexor BCD being substantially longer than other muscles. The average fascicle lengths of the anti-gravity shoulder protractors and retractors were 27.2 ± 5.6 mm and 30.5 ± 15.0 mm, respectively (Fig. 3C). Fascicle lengths of elbow flexors (46.6 ± 30.5 mm) exceeded the lengths of shoulder adductors/adductors, forearm supinator/pronator, and wrist plantarflexors/ abductors (p < 0.005) but were not significantly different from elbow extensors (p = 0.06; 28.7 ± 8.2 mm). Fascicle length of antigravity wrist plantarflexors and their antagonists wrist dorsiflexors were not significantly different either (12.7 ± 6.0 mm vs. 20.5 ± 13.8 mm, p = 0.07). This contrasts with the hindlimb, where antigravity muscles (quadriceps, triceps surae) have substantially shorter fascicles than their counterparts (hamstrings, tibialis anterior) (Sacks and Roy, 1982).

### Forelimb locomotor mechanics

Forelimb kinematic and kinetic variables obtained in all animals during overground locomotion are shown as a function of the normalized cycle time in Fig. 4. The mean shoulder, elbow, and wrist angles (see Fig. 4A, B, C) showed patterns and joint angle ranges typical for cat walking (Miller and Van Der Meche, 1975; English, 1978b; Lavoie et al., 1995; Prilutsky et al., 2005). During the swing phase, shoulder and elbow angles reached their minimum values (86.8 ± 6.6° and 78.5 ± 7.3°) and the wrist its maximum value (232.7 ± 15.8°) in early to mid-swing. These instances correspond to maximal shoulder retraction, elbow flexion, and wrist dorsiflexion. The maximum shoulder protraction (124.3 ± 4.5°) and elbow extension (113.5 ± 7.5°) occurred at the end of swing. During the stance phase, the shoulder retracts, while the elbow joint continues extending. The wrist joint is slightly dorsiflexing in stance before plantarflexing at end-stance.

**Figure 4.**
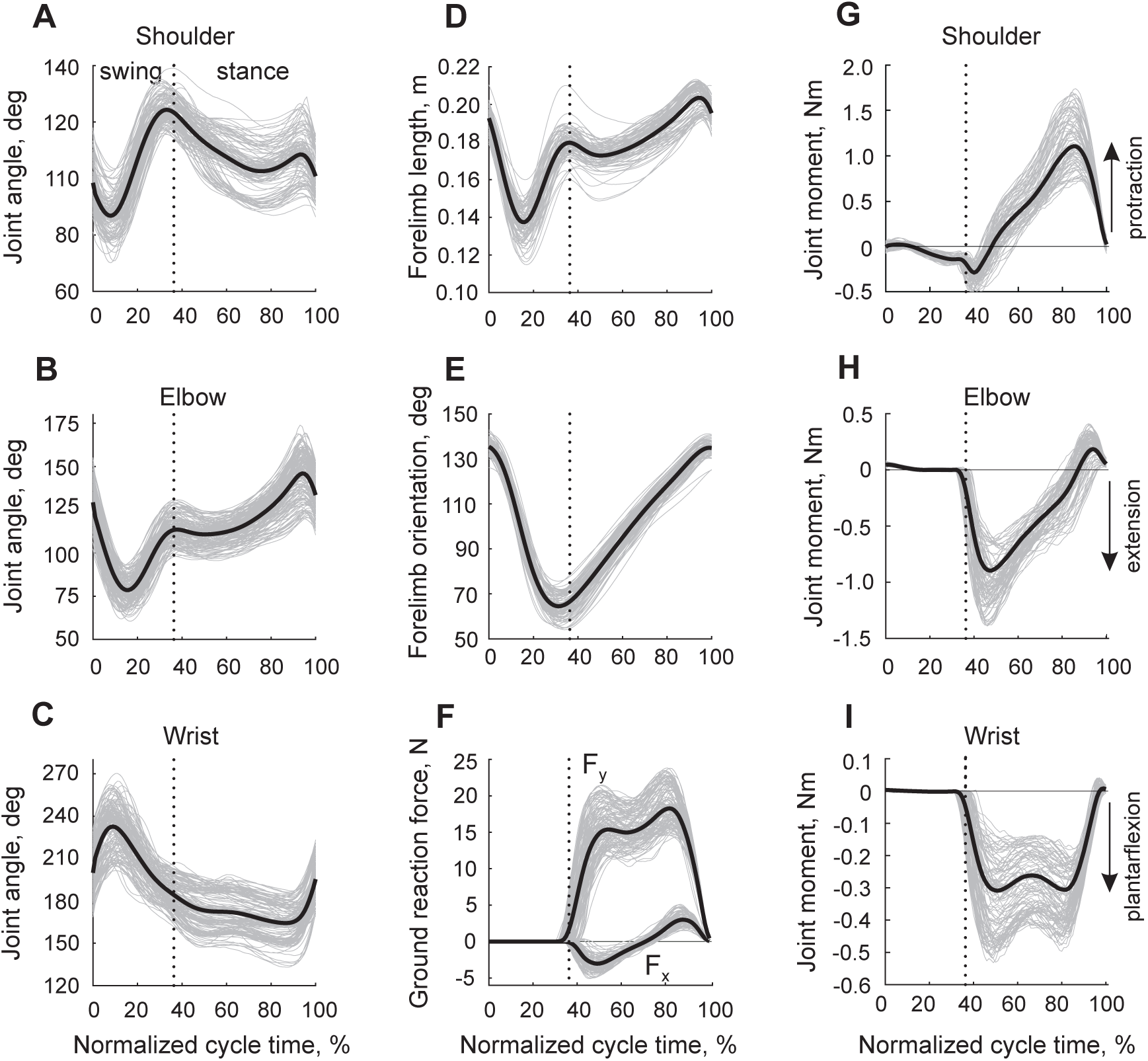
Forelimb mechanical variables during the cycle of overground locomotion (mean patterns are shown by thick black lines, individual cycles of different animals are shown by thin gray lines). The mean patterns were obtained from all studied cats (n=7) and walking cycles (n=211), see Table 2. **A**, **B**, **C**: Joint angles at the shoulder, elbow, and wrist, respectively. **D**, **E**: Forelimb length and forelimb orientation, respectively, as defined in Fig. 1A. **F**: Vertical (F_y_) and horizontal (F_x_) components of the ground reaction force vector. **G**, **H**, **I**: Resultant moments of force at the shoulder, elbow, and wrist joints. Vertical dotted lines separate the swing and stance phase.

The pattern of the forelimb length (the distance between the shoulder and the paw, Fig. 1A, sagittal view) closely resembled the elbow angle pattern with the shortest length (0.137±0.009 m) in mid-swing and the peak value (0.204±0.006 m) at the end of stance (Fig. 4D). The forelimb orientation changes from its maximum value (136.4±2.8°) at swing onset to its minimum value (64.2±5.0°) at the end of swing (Fig. 4E). The patterns of forelimb length and orientation were remarkably similar to those of the hindlimb during walking in the cat (Chang et al., 2009; Klishko et al., 2014).

The patterns and peak values of the ground reaction forces exerted by the forelimb on the ground (Fig 4F) were typical for cat walking (Lavoie et al., 1995; Farrell et al., 2014). Specifically, the vertical force peak (19.0 ± 2.6 N or 58.1 ± 8.0% body weight) occurred near the end of stance. Note that this value is approximately 20% higher than the vertical peak force of the cat hindlimb, which occurs in early stance during walking at comparable speeds (Lavoie et al., 1995; Farrell et al., 2014). The horizontal component of the ground reaction force had a negative peak in the first half of stance (−3.2 ± 1.0 N) and a positive peak in the second half (3.2 ± 0.9 N). Although the absolute values of the positive and negative peaks were the same, the area under the negative forces (force impulse) was larger, indicating that the net action of the forelimb during stance is to decelerate the body, as opposed to the hindlimbs that accelerate the body during stance (Lavoie et al., 1995; Farrell et al., 2014).

The resultant joint moments of force characterizing the action of forelimb muscles in the sagittal plane are shown in Fig. 4G-I. Wrist plantarflexors and elbow extensors produced their largest moment during stance (peak values were −0.339 ± 0.113 Nm and −0.998 ± 0.225 Nm, respectively; Fig. 4 H, I). There are relatively small wrist dorsiflexion and elbow flexion moments at the end of stance and early swing, and the wrist dorsiflexors and elbow flexors continue producing flexor moments during the first half of swing, although their peak flexor moment values are very low (0.004 ± 0.002 Nm and 0.052 ± 0.010 Nm, respectively). The action of shoulder protractors located in front of the shoulder joint produced their moment of force throughout most of the stance phase with a maximum protraction moment of 1.181 ± 0.283 Nm appearing closer to stance offset (Fig. 4G). The peak shoulder retraction moment (−0.305 ± 0.091 Nm) occurred at stance onset. During the first third of swing, the shoulder moment is protraction (the peak of 0.023 ± 0.028 Nm) before switching to retraction for the rest of swing. Patterns of wrist and elbow moments resembled corresponding ankle and knee moments of walking cats, although the magnitude of the elbow extension moment was over two times greater than the peak knee extension moment (McFadyen et al., 1999; Gregor et al., 2018). The shoulder protraction moment occurred during most of stance, as opposed to the corresponding hip flexion moment, which acts during the second half of the stance phase (McFadyen et al., 1999; Gregor et al., 2018).

The forelimb MTU lengths normalized by the optimal MTU length (see Table 5) are shown as a function of the normalized cycle time in Fig. 5A. Length changes of one-joint muscles were closely related to the joint angle patterns. The shoulder retractors located posterior to the shoulder joint increased their length from a minimum in early swing to a peak value before swing offset. The antagonists, shoulder protractors, had opposite length changes with peak length in early swing and minimum length just before stance onset. MTU length changes of the one-joint elbow flexors followed the pattern of the elbow joint angle with minimum length in mid-swing and two local length peaks just before stance onset and stance offset. MTU length changes of one-joint elbow extensors were opposite. Patterns of MTU length changes of wrist dorsiflexors and plantarflexors were also opposite to each other with the pattern of wrist dorsiflexors demonstrating maximum length in early swing and a relatively small decrease in length during stance. The two-joint antagonists BB and TLONG likewise demonstrated opposite length changes with BB reaching a minimum MTU length in mid-swing and peak length at stance offset. The MTU length of the elbow flexors-wrist dorsiflexors similarly had a minimum in mid-swing and maximum at stance offset. The patterns of forelimb MTU length changes were generally consistent with those of cat hindlimb MTUs (Goslow et al., 1973; Gregor et al., 2006; Klishko et al., 2021). Specifically, proximal forelimb flexors (BB) and hindlimb flexors (iliopsoas, rectus femoris, sartorius medial head) located anterior to the shoulder or hip joints reach their maximum length at stance offset. The proximal antagonists, shoulder retractors and hip extensors reach their peak MTU lengths prior to stance onset. There were also differences in the MTU length patterns between the forelimb and hindlimb, as there is a much less pronounced stretch of the forelimb distal extensors-plantarflexors in early stance due to smaller yield at the wrist and elbow compared to the stretch of ankle plantarflexors and knee extensors.

**Figure 5.**
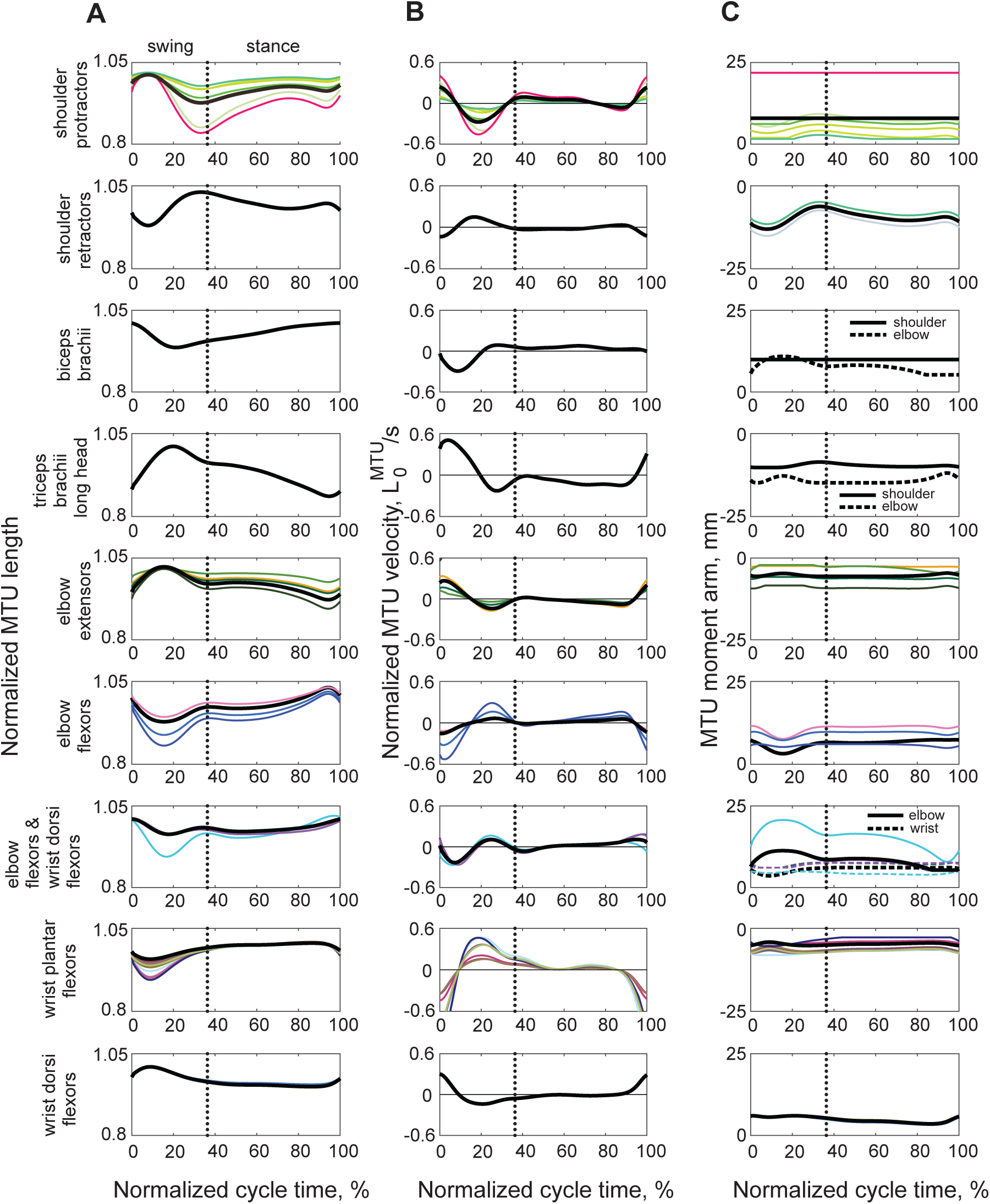
Computed mean normalized length (**A**), normalized velocity (**B**) and moment arms (**C**) of individual MTUs (thin color lines) and the equivalent MTUs representing individual MTUs with the same action at joints during the walking cycle. Colors of individual MTU patterns correspond to the colors of individual MTUs in Fig. 1A. MTU length is normalized to the optimal MTU length (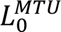, Table 5); MTU normalized velocity units are 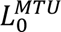/*s*, where 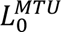 is the optimal MTU length (Table 5). Positive moment arm corresponds to the direction of MTU moment of force that tends to rotate the distal link counterclockwise (see the moment sign convention in Fig. 1). The mean patterns were computed from all studied cats (n=7) and walking cycles (n=211), see Table 2.

The normalized MTU velocity changes during the walking cycle were generally similar across forelimb muscles (Fig. 5B). The greatest values of MTU shortening and lengthening occurred in swing, and relatively constant velocities were seen during most of stance. Peak lengthening velocities, which are related to high activity of spindle group Ia afferents (see below), often occurred in proximal MTUs at the stance-swing transition (shoulder protractors, triceps brachii long head, one-joint elbow extensors), and in distal wrist dorsiflexors.

The forelimb moment arms changed relatively little during the walking cycle for most MTUs, except for shoulder retractors (range: 7.4 – 15.1 mm for TJ and 4.9 – 11.5 mm for SD) and ECR (range: 7.9 – 20.7 mm); Fig. 5C. The absolute value of the moment arm at a joint determines the range of MTU length change and peak MTU velocity (e.g., compare the moment arms, lengths, and velocity peaks among the shoulder protractors or elbow extensors; Fig. 5). As evident from Fig. 5C, moment arm values of proximal MTUs, such as shoulder protractors and retractors, BB, and TLONG, were substantially longer than those of distal MTUs of wrist dorsiflexors and plantarflexors.

### Maximum moments of individual and equivalent MTUs during locomotion

The k-means clustering algorithm applied to the mean peaks of maximum moments of force produced by 40 MTUs during walking of all cats revealed 9 clusters or MTU groups with unique muscle actions at forelimb joints in the sagittal plane (Fig. 1B). These included shoulder protractors (6 MTUs), shoulder retractors (2 MTUs), shoulder protractor-elbow flexor (1 MTU), shoulder retractor-elbow extensor (1 MTU), elbow extensors (4 MTUs), elbow flexors (3 MTUs), elbow flexors-wrist dorsiflexors (5 MTUs), wrist plantarflexors (16 MTUs), and wrist dorsiflexors (2 MTUs). Because synergistic muscles are normally activated together (Buchanan et al., 1986; Prilutsky, 2000b, a; Hug et al., 2022), it is possible to simplify the forelimb musculoskeletal model by substituting the action of individual MTUs in each cluster group by an equivalent MTU with a similar muscle action. The morphological and geometric parameters of such equivalent muscles were obtained as described in the Methods and are listed in Table 5 (bold font) and Table 7. The comparison of the sum of the maximum moments of each cluster with the corresponding moment of the equivalent MTU, computed assuming maximum activation of the MTUs, *A* = 1 (see Eqs. 2-6), demonstrated a close match (Fig. 6A-I). The normalized maximum difference did not exceed 7% for all MTU clusters except for one (elbow flexors, 11%).

**Figure 6.**
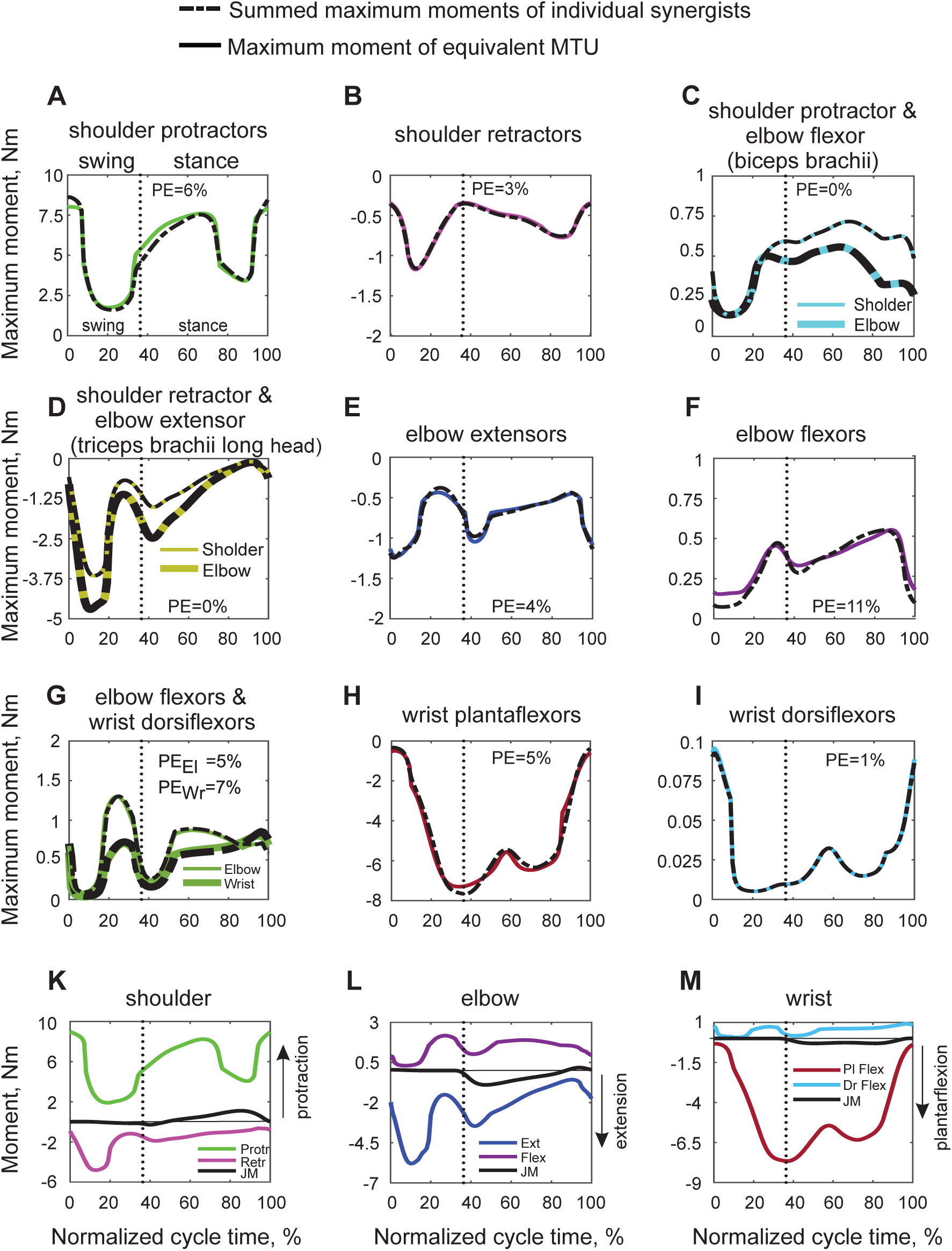
Mean maximum moments of force and mean resultant moments of force at forelimb joints during the walking cycle. The vertical dotted lines separate the swing and stance phase. The moment sign convention is the same as in Fig. 1A, i.e. positive direction corresponds to the counterclockwise rotation of the distal segment at the joint. The mean patterns were computed from all studied cats (n=7) and walking cycles (n=211), see Table 2. **A – I**: Comparisons of the summed maximum moment of force produced by individual synergists at joints (black dashed lines) and by the equivalent MTU with the same mechanical action at the joints (continuous color lines). Colors of moments produced by the equivalent MTUs correspond to the colors of equivalent muscles in Fig. 1B. Two-joint MTUs (biceps brachii, triceps brachii long head, and elbow flexors-wrist dorsiflexors) produce maximal moments at each joint they span. PE is the percent error between the sum of maximum moments produced by synergists at the joints and by their equivalent muscle. **K – M**: Comparisons of the maximum moments of force produced by all equivalent MTUs in both directions with the actual mean moment of force (see Fig. 4G-I) during the walking cycle. Protr and Rets, protraction and retraction; Ext and Flex, extension and flexion; Pl and Dr, plantar and dorsi.

The maximum potential for moment production occurred in the antigravity muscle groups (shoulder protractors, elbow extensors, and wrist plantarflexors) during stance or at the stance-swing transition. Their antagonists demonstrated peak maximum moments during swing (shoulder retractors, shoulder protractor-elbow flexor BB, shoulder retractor-elbow extensor TLONG, and wrist dorsiflexors). The two-joint MTUs contributed greater maximum moments of force during stance to the joints in which these MTUs produce an extension action, such as shoulder protraction for BB (Fig. 6C) and elbow extension for TLONG (Fig. 6D). The elbow flexors-wrist dorsiflexors produced substantially greater peaks of maximum moment for elbow flexion during the second half of swing and the middle portion of stance (Fig. 6G).

The resultant moments of force during walking determined by inverse dynamics analysis were substantially lower in magnitude than the maximum possible moments, except for elbow and wrist dorsiflexion moments (Fig. 6K-M). The greatest unused moment potential occurred in wrist plantarflexors, which could be related to the additional functions of the forelimb, including hunting and defense.

### Motor output of forelimb MTUs during walking

We computed activation of each MTU during the walking cycle by minimizing the cost function of minimum fatigue (eq. 16) under the constraints of muscle contractile force-length-velocity properties (eqs. 2-6), resultant joint moments of force during walking (eq. 17), and minimum and maximum activations (eq. 18). Antigravity individual MTUs and equivalent MTUs representing the functional groups revealed by the cluster analysis were activated primarily during the stance phase (Fig. 7A; shoulder protractors, elbow extensors, wrist plantarflexors). Their antagonists (shoulder retractors, elbow flexors, two-joint elbow flexors-wrist dorsiflexors, but not one-joint wrist dorsiflexors) were mostly active during swing. The two-joint shoulder protractor-elbow flexor BB was active at end stance and early swing, whereas the two-joint shoulder retractor-elbow extensor TLONG was active from end swing and throughout most of stance. No activation was predicted for one-joint wrist dorsiflexors (Fig. 7A), although the resultant wrist dorsiflexion moment of force was flexor in early swing and late stance (Fig 4 I). All synergistic MTUs within each functional group were activated with generally similar patterns, except for pronator teres (PT) among elbow flexors. In 3D, PT is primarily a pronator and has extremely short fascicles, relative to other elbow flexors. The computed activations of the one-joint antagonists (shoulder protractors-retractors and elbow extensors-flexors) demonstrated reciprocal activation. Two-joint muscles often had peak activation when the moments produced by these muscles at both joints coincided with the direction of the resultant joint moments. For example, BB had two activity bursts, in early swing and late stance (Fig. 7A), when there was simultaneous production of resultant shoulder protraction moment (Fig. 4G) and resultant elbow flexion moment (Fig. 4H). The TLONG initiated its activity burst earlier than the one-joint elbow extensors (in late swing and swing-stance transition, Fig. 7A) when there was a combination of shoulder retraction (Fig. 4G) and elbow extension resultant moments (Fig. 4H). Two of the three activation peaks of two-joint elbow flexors-wrist dorsiflexors at the end of stance and early swing transition (Fig. 7A) coincided with the simultaneous production of a resultant flexion moment at the elbow (Fig. 4H) and a dorsiflexion moment at the wrist (Fig. 4 I).

**Figure 7.**
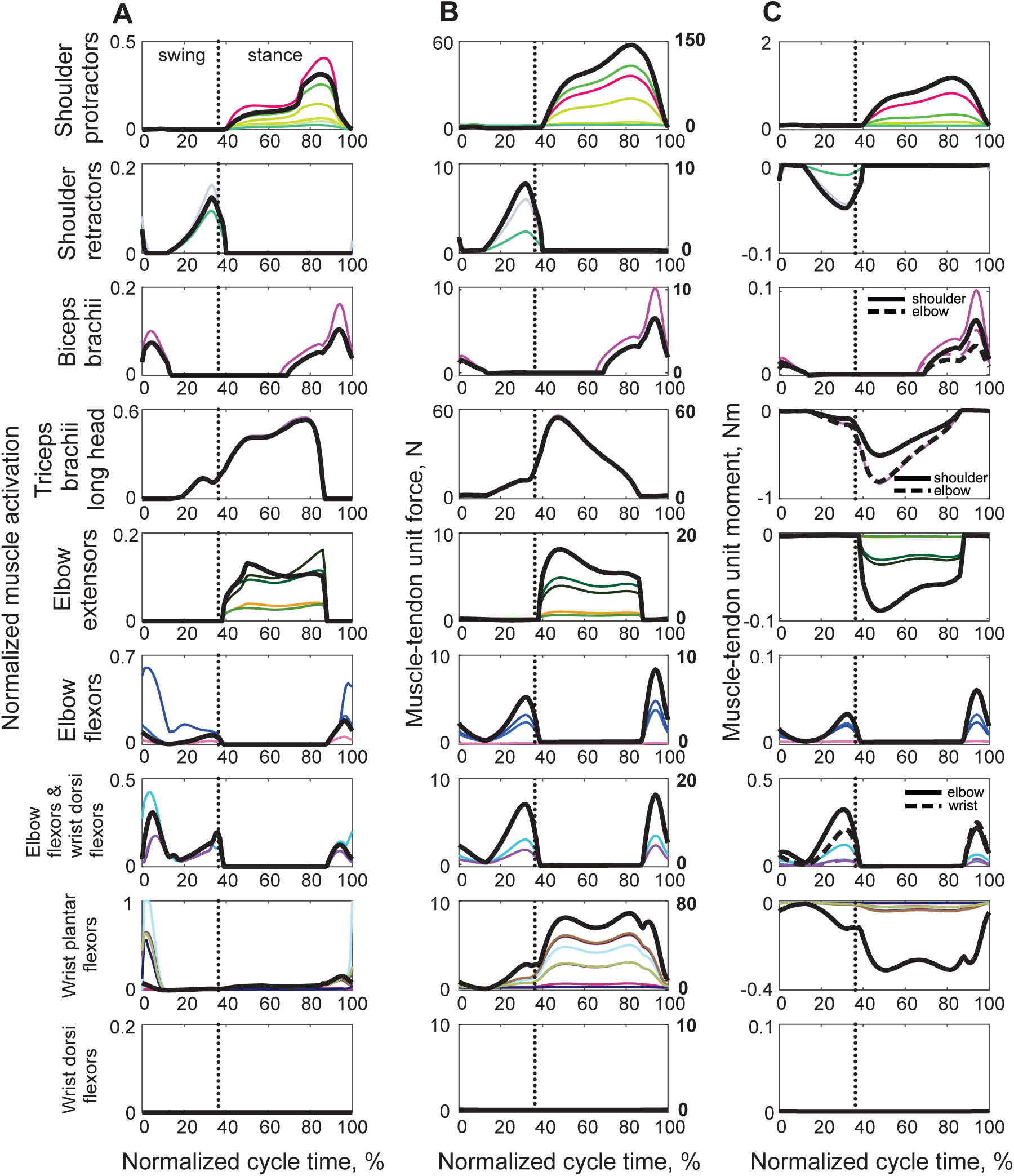
Mean computed activation (**A**), force (**B**), and moment of force (**C**) of individual MTUs (thin color lines) and the equivalent MTUs (thick black lines) during the walking cycle. Two-joint MTUs produce moments of force at each joint they span. The vertical dotted lines separate the swing and stance phase. The moment sign convention is the same as in Fig. 1A. Colors of individual MTU patterns correspond to the colors of individual MTUs in Fig. 1A. The mean patterns were computed from all studied cats (n=7) and walking cycles (n=211), see Table 2.

The computed activation when compared with the recorded EMG activity of selected muscles showed that such minimum fatigue minimization generally predicted the reciprocal activation of antagonist forelimb muscles, such as TLAT vs PT and BR, and BB vs TLONG, as well as synergistic activation of agonists, such as TLONG with TLAT and PT with BR (Fig. 8). Computed activations using the models with 40 MTUs and 9 equivalent MTUs were generally similar except for PT where the 40-MTU model demonstrated a closer match with the recorded EMG activity (Fig. 8). Both models showed that the computed activity patterns of SPS and both heads of triceps brachii were qualitatively similar to EMG activity (Fig. 8). For the other muscles, the computed activations by the two models showed some discrepancies with EMG activity (Fig. 8).

**Figure 8.**
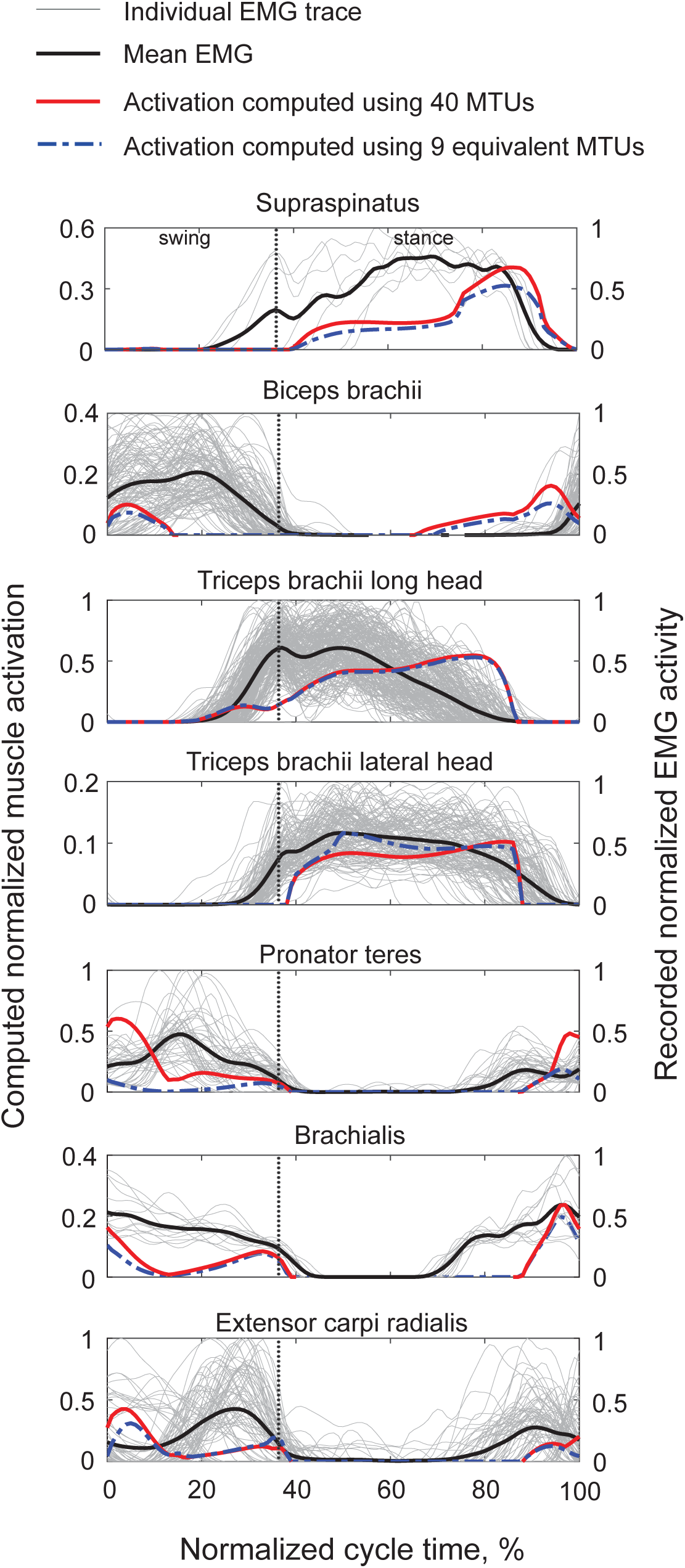
Comparison of mean patterns of computed activation of selected MTUs with the recorded EMG activity. The vertical dotted lines separate the swing and stance phase. The computed activation has the range from 0 to 1. The recorded EMG patterns were normalized by the peak value across all cycles within each animal. The computed mean activation patterns were obtained from all studied cats (n=7) and walking cycles (n=211), see Table 2. Red lines show activations computed using the full forelimb musculoskeletal model with 40 MTUs; blue lines show activations computed using a reduced forelimb model with 9 equivalent muscles (see eqs. 16-18 and Fig. 1B). The mean patterns of recorded EMG (thick black lines) are averages of 531 cycles (thin gray lines) of 16 cats (Table 4).

Computed MTU forces generally had similar patterns to the computed activations with some minor exceptions (Fig. 7B). This is expected given that MTU force also depends on MTU length and velocity of length change (see eqs. 2-6). This also explains the difference in the magnitudes of the activation and forces for individual MTUs within functional groups (e.g. shoulder protractors and retractors). The computed forces of the functional groups were much larger for proximal antigravity muscles, shoulder protractors and TLONG, with peak forces approaching 60 N. The peak forces of other MTUs did not exceed 10 N.

Patterns of the moments of force of individual and equivalent MTUs resembled the force patterns of the same MTUs (Fig. 7C), indicating relatively small changes in the MTU moment arms during the walking cycle. The moments produced by the two-joint muscles at the joints they span had different magnitudes; protraction shoulder moment of BB was greater than flexion elbow moment, elbow extension moment of TLONG exceeded shoulder retraction moment, and elbow flexion moments of elbow flexors-wrist dorsiflexors were greater than wrist dorsiflexion moments.

### Computed proprioceptive activity of forelimb MTUs

The developed forelimb musculoskeletal model allowed computing the activity of muscle spindle group Ia and II afferents using the calculated velocity, length (Fig. 5A,B), and activation (Fig. 7A) of individual MTUs (eqs. 7, 8), as well as the activity of Golgi tendon organ group Ib afferents using the calculated MTU forces (Fig. 7B; eq. 9). Activity patterns of Ia afferents (Fig. 9A) resembled patterns of MTU velocities. Specifically, one-joint shoulder protractors, two-joint TLONG, one-joint elbow extensors, and one-joint wrist dorsiflexors reached their peak Ia activity at the stance-swing transition, which corresponded to the peak velocities of these MTUs (Fig. 5B). One-joint shoulder retractors, two-joint BB, one-joint elbow flexors and wrist plantarflexors reached their peak Ia activity between the middle and the end of swing.

**Figure 9.**
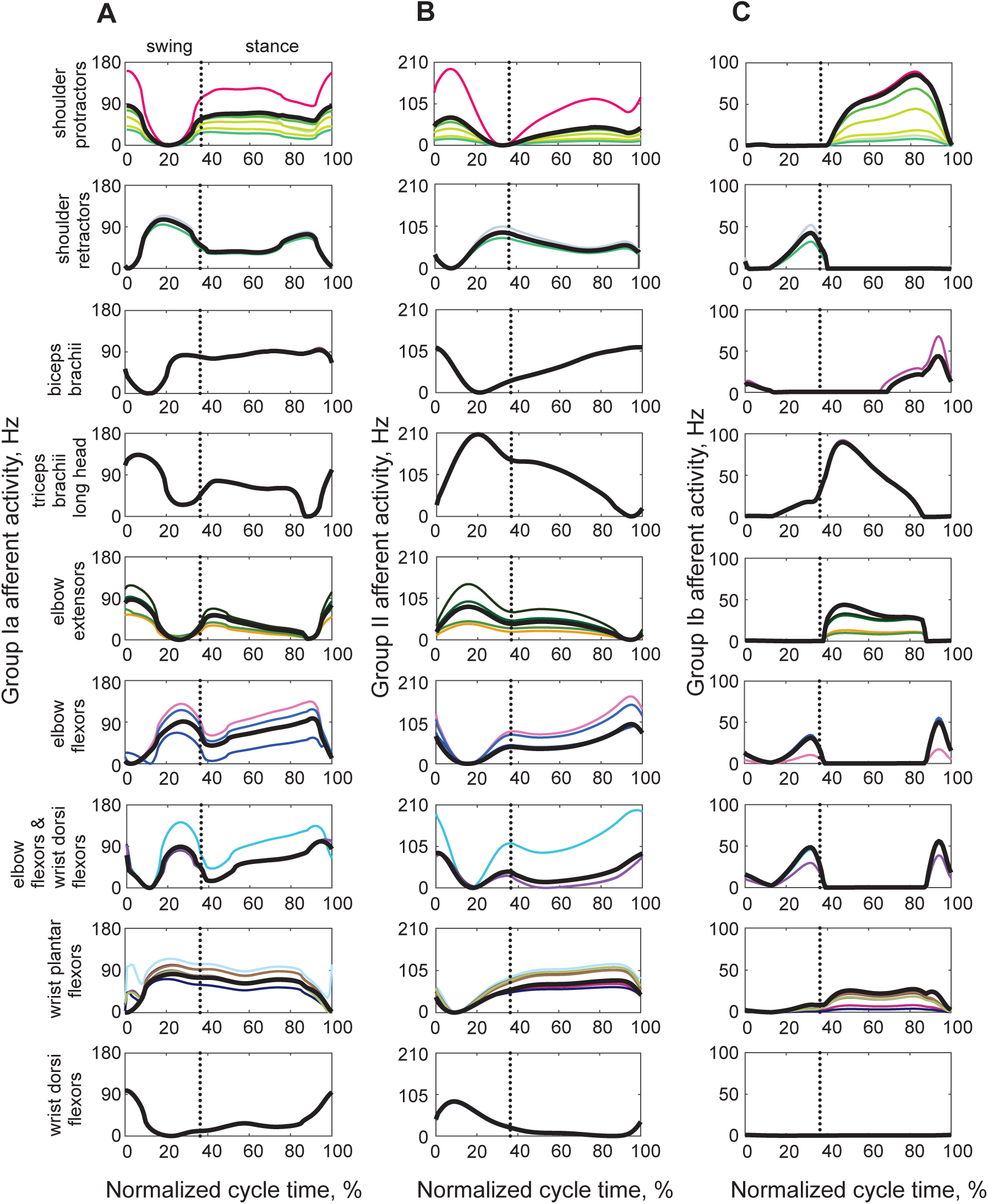
Mean computed activity of muscle spindle group Ia afferents (**A**), muscle spindle group II afferents (**B**), and Golgi tendon group Ib afferents (**C**) of individual MTUs (thin color lines) and of the equivalent MTUs (thick black lines) during the walking cycle. The vertical dotted lines separate the swing and stance phase. Colors of individual MTU patterns correspond to the colors of individual MTUs in Fig. 1A. The mean patterns were computed from all studied cats (n=7) and walking cycles (n=211), see Table 2.

The muscle spindle group II activity closely matched length changes of the corresponding MTUs (Figs. 5A, 9B). MTUs located anteriorly to the forelimb joints (one-joint shoulder protractors, two-joint BB, one-joint elbow flexors and two-joint elbow flexors-wrist dorsiflexors) reached their maximum length and spindle group II activity near the stance-swing transition. One-joint shoulder retractors and wrist plantarflexors reached their maximum length and spindle group II activity at the swing-stance transition. The magnitude of MTU length- and velocity-dependent afferent activities of individual MTUs increased with the MTU moment arm, the distance between the line of MTU action and the joint center. For example, the shoulder protractor SPS (pink trace in Figs. 1A, 9A,B) was farthest from the shoulder center and demonstrated the highest rates of spindle activity. This is because MTU length change due to joint angle changes is proportional to the MTU moment arm (An et al., 1983).

Because the firing rate of Ib afferents is proportional to muscle force (see eq. 9), the patterns of MTU muscle forces and group Ib afferent activities were identical (Figs. 7B, 9C). Thus, antigravity forelimb muscle group Ib afferents were active during stance, while Ib afferent activity of their antagonists occurred during the swing phase.

The peak rates of computed proprioceptive activity did not exceed 200 Hz, which is consistent with the rates of proprioceptive afferents of hindlimb muscles during cat locomotion (Loeb and Duysens, 1979; Prochazka and Gorassini, 1998). Our computed rates were also consistent with rates of group Ia afferents of selected shoulder protractors and retractors and elbow flexors and extensors maintained at a constant length while the other three limbs of the cat decerebrate preparation performed treadmill locomotion (Cabelguen et al., 1984).

We used the computed activation and proprioceptive activity of 40 MTUs during walking, the known distribution of motoneuron pools of selected cat forelimb muscles in cervical segments, Table 8 (Fritz et al., 1986a; Fritz et al., 1986b; Horner and Kummel, 1993), and assumed the same rostro-caudal distribution of motoneuron pools and their proprioceptive inputs in the cervical spinal cord (Sterling and Kuypers, 1967b, a; Levine et al., 2012) to compute spatiotemporal maps of motor and sensory activity in cervical spinal segments during the walking cycle (Fig. 10).

**Figure 10.**
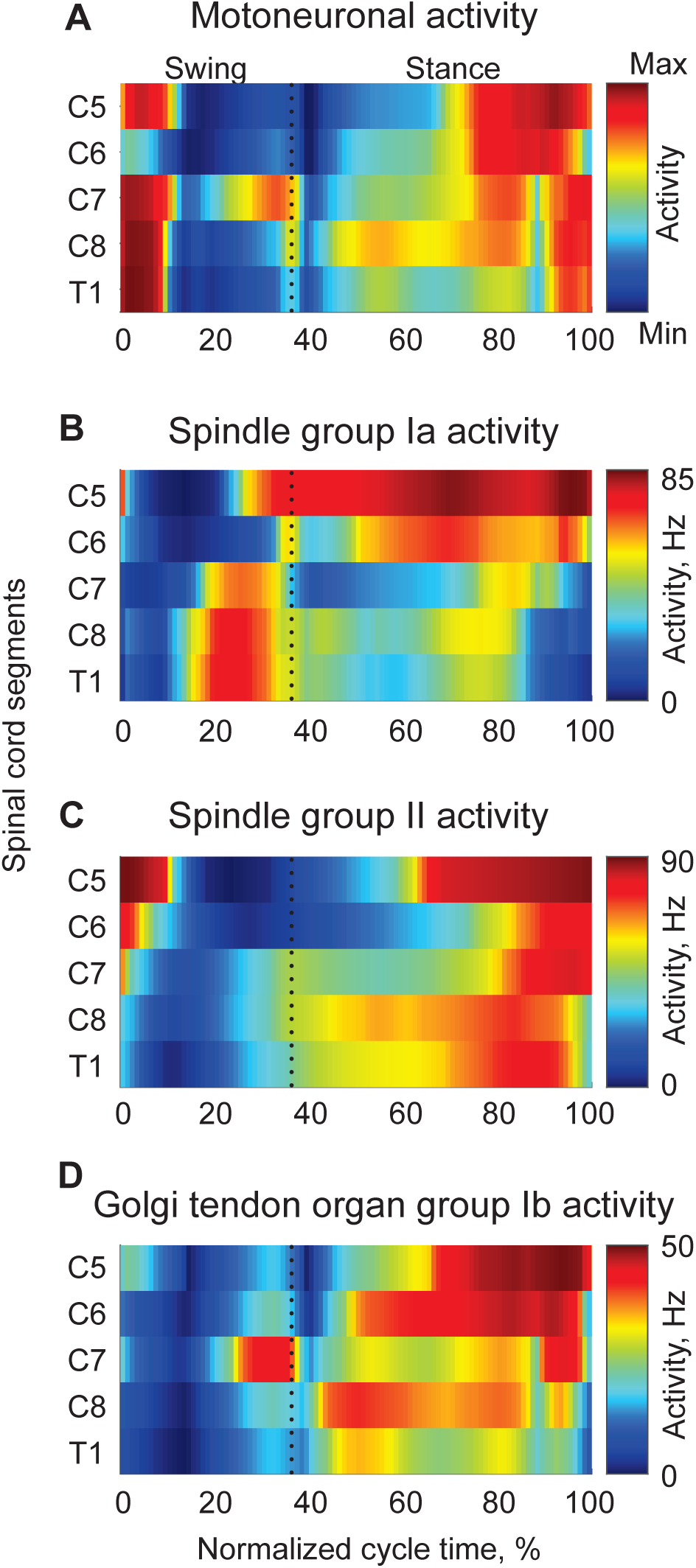
Spatiotemporal maps of computed motoneuronal activation (**A**), and afferent activity of spindle group Ia (**B**), spindle group II (**C**), and Golgi tendon organ group Ib (**D**) afferents projected to the corresponding cervical spinal segments during the walking cycle (see text and eq. 7-9 and 19 for details). The vertical dotted lines separate the swing and stance phase. The spatiotemporal maps were computed from all studied cats (n=7) and walking cycles (n=211), see Table 2.

According to Table 8, segment C5 contains a relatively small proportion of motoneurons (1-9%) of one-joint shoulder protractors, two-joint elbow flexors-wrist dorsiflexors (9%), and one-joint wrist dorsiflexors (3%). Motoneurons of the two-joint BB constituted a substantial proportion of the total at C5 (37%). Segments C6 and C7 contained the largest proportions of motoneurons innervating mostly proximal muscles: the one-joint shoulder protractors (between 20% and 87% for most MTUs), one-joint retractors (33-67% for SD, 1-99% for TJ), the two-joint BB (19-40%), one-joint elbow flexors (1-94%), and the two-joint elbow flexor-wrist dorsiflexor ECR (32-59%). Segments C8 and T1 contained the largest proportion of motoneurons of the two-joint elbow flexor-wrist dorsiflexor EDC (63% and 36%) and one-joint wrist plantarflexors (28-72%). Since we could not find the distribution of motoneurons in the two-joint MTUs spanning the shoulder and elbow (BB and TLONG) and one-joint elbow extensors, we took these data from the corresponding MTUs of monkey (Jenny and Inukai, 1983). Most BB motoneurons in the monkey’s spinal cord are located in segments C5 – C7 (19-37%), while TLONG and elbow extensors are in segments C7 – T1 (22-52%). In that monkey study, the authors compared the distribution of motoneurons of the EDC between 4 monkeys and 3 cats and found that the total number of motoneurons was 295.0 ± 30.2 and 299.0 ± 19.2, respectively, and all of them were in segments C7, C8, and T1. The distribution of motoneurons for these segments was 1.1 ± 1.0%, 55.0 ± 7.2%, 43.9 ± 7.9% for the monkeys and 2.3 ± 2.1%, 75.8 ± 20.6%, 21.3-23.6% for the cats, respectively. The mean motoneuron distributions of EDC from (Jenny and Inukai, 1983) were within one standard deviation of both the monkey EDC and the cat EDC data of (Fritz et al., 1986b). Thus, substituting the missing cat data for the three MTU groups in Table 8 with data from the monkey is reasonable.

The computed spatiotemporal maps of normalized motoneuronal activity shown in Fig. 10A indicate the total proportion of motoneurons recruited in segments C5-T1 during the overground walking cycle. Thus, MTUs with high normalized activity during walking, like some one-joint wrist plantarflexors and elbow flexors in early swing (Fig. 7A), had the greatest contribution to the spatiotemporal map. The normalized motoneuronal activity was high during the stance phase, the stance-swing transition and early swing, and it was distributed across segments C5 through T1 (Fig. 10A). An elevated activity in the stance phase started at the swing-stance transition and occurred mostly in segments C7 and C8 because of muscles involved in shoulder retraction with simultaneous elbow extension and simultaneous elbow flexion and wrist dorsiflexion (see also Fig. 7A and Table 8). The stance-related activity of motoneurons shifts from caudal segments C7-T1 in early to mid-stance to segments C5 – C7 in the second part of stance from ∼70% to 90% of the cycle. The highest proportion of active motoneurons during late stance occurred in rostral segments C5 and C6. This activity reflected recruitment of shoulder protractors and two-joint biceps brachii. Motoneurons of elbow extensors and wrist plantarflexors, which are mostly located in more caudal cervical segments, were also recruited during late stance but to a lower degree. At end stance (90-100% of the cycle) and early swing (0-10%), relatively high motoneuronal activity occurred in segments C5-T1 that was related to recruitment of the two-joint shoulder protractor-elbow flexor BB and elbow flexors-wrist dorsiflexors (ECR, EDC), as well as one-joint elbow flexors and wrist plantarflexors (Fig. 7A). During most of the swing phase, shoulder retractors, elbow flexors and wrist dorsiflexors were active, but a relatively small proportion of their motoneurons was recruited across segments C5-T1 (Fig. 10A).

The spatiotemporal map of spindle group Ia afferent activity that serves as sensory input to the motoneurons in the corresponding spinal segments is shown in Fig. 10B. Group Ia activity started increasing from its minimum level at mid-swing. This activity was initiated at caudal segments T1 and C8 (afferents of one-joint wrist plantarflexors and two-joint elbow flexor-wrist dorsiflexor EDC, Fig. 9A) and continued rostrally to segments C7 (two-joint elbow flexor-wrist dorsiflexor ECR, one-joint elbow flexors, two-joint elbow flexor-shoulder protractor BB, one-joint shoulder retractors) and C5 (BB). During the stance phase, the greatest Ia activity occurred in segments C6 and C5 (afferents of one-joint shoulder protractors and elbow flexors, two-joint BB and ECR).

The spatiotemporal map of spindle group II afferent activity started increasing at end swing and the swing-stance transition, especially in segments C7 and C8, associated with increasing length of one-joint shoulder retractors, elbow flexors, and wrist plantarflexors, as well as two-joint elbow flexors-wrist dorsiflexors (Figs. 9B and 10C). This activity became progressively greater during stance in segments T1 and C8 (wrist plantarflexors and two-joint EDC) and C5 (two-joint BB). The peak of group II afferent activity occurred at the stance-swing transition in segments C5 – C7. This activity corresponded to recruitment of group II afferents of two-joint BB and ECR and one-joint elbow flexors and shoulder protractors (Figs. 9B and 10C).

The spatiotemporal map of group Ib afferent activity was similar to the map of motoneuronal activity. All segments from C5 to T1 had higher activity during stance and this activity was associated with one-joint shoulder protractors, elbow extensors and wrist plantarflexors, as well as two-joint BB (Fig. 10D). One noticeable difference between the spatiotemporal maps of motoneuron and Ib activation was the lack of high Ib activity in early swing because of low force production by wrist plantarflexors and elbow flexors despite their relatively high computed motoneuronal recruitment.

## DISCUSSION

The goal of this study was to determine morphological characteristics of the cat’s forelimb muscles and evaluate their contributions to kinetics of locomotion and motion-dependent sensory feedback. We addressed this goal by measuring forelimb morphological characteristics post-mortem and by recording forelimb kinematics, ground reaction forces, and EMG activity of selected muscles during walking. We also used the computed length, velocity, activation, and force of each forelimb MTU during walking to estimate the motion-dependent sensory activity of muscle proprioceptors. Such comprehensive information allows us to analyze the role of forelimb morphology in mammalian locomotor with the level of detail unavailable previously.

Before discussing potential contributions of forelimb morphology to motor output and sensory feedback during locomotion, we consider the effects of potential errors in determining morphological characteristics and some assumptions on computed variables.

### Effects of model sensitivity to model parameter changes and assumptions

As discussed in the Methods, the computed motor outputs and proprioceptive signals of the forelimb model depend on multiple model parameters, including optimal MTU length, maximal muscle shortening velocity, PCSA, pennation angle (see eqs. 1-6, Table 5), and geometric parameters describing the MTU path with respect to the joints (Fig. 2, Table 7). To evaluate sensitivity of the computed maximum moments of each equivalent MTU averaged over the walking cycle to changes in major morphological and geometric MTU parameters, we computed the first-order global sensitivity index for each parameter based on the Sobol method (Sobol, 2001) implemented in the Global Sensitivity Analysis Toolbox (flax, 2022). The first-order indices (main effects) measure the contribution of each model parameter to the total output variance (the variance of the mean maximum moment in our case), neglecting interaction effects among the parameters; the first-order index has a range of 0 to 1 (Qian and Mahdi, 2020). The effects of random changes in each parameter within 10% of its nominal value from Tables 5 and 7 on the computed first-order sensitivity indices are shown in Fig. 11. The maximum moment is most sensitive to changes in PCSA for all muscle groups, followed by geometric parameters defining the locations of the MTU origin and insertion that in turn define the MTU moment arms. The optimal muscle fascicle length affects the variance of the maximum moment of the two-joint shoulder retractors-elbow extensors, wrist dorsiflexors, and shoulder protractors, although to a lesser extent. The normalized maximum muscle shortening velocity, percentage of slow-twitch fibers, and pennation angle do not have noticeable effects on the maximum moment variance. Previous musculoskeletal modeling studies likewise found that the resultant moment of force at the joint and related mechanical variables are most sensitive to changes in PCSA and muscle moment arms (Raikova and Prilutsky, 2001; Ramalingasetty et al., 2021). The PCSA can be accurately obtained postmortem using eq. 1. The situation is more complex for determining accurate locations of the MTU origin and insertion sites because of their broad and complex 3D areas in most muscles. The moment arm value is also affected by the accuracy of determining the joint rotation axis or center. One way to improve the accuracy of determining the MTU moment arm is to use the tendon-excursion method (An et al., 1983), which does not require the muscle attachment points and the joint center of rotation. If coordinates of MTU attachment points are required to compute MTU length and velocity during movement, the accuracy determining these and other parameters can be improved by adjusting the measured parameters by minimizing the difference between the recorded and simulated movement mechanics and muscle activity (Neptune et al., 2008; Bunderson et al., 2012; Prilutsky et al., 2016).

**Figure 11.**
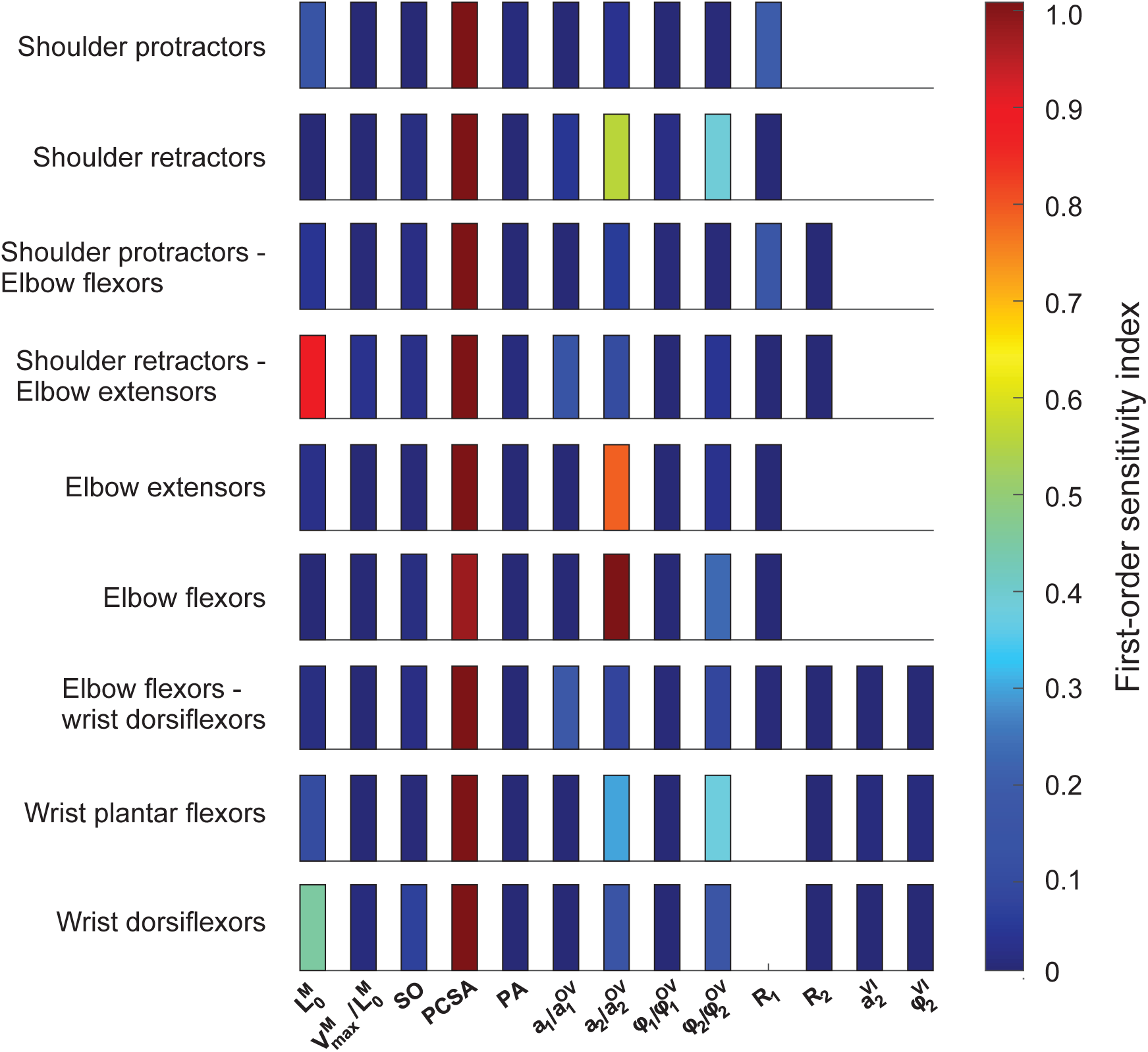
Sensitivity of the computed maximum moment of force averaged over the walking cycle for each equivalent MTU to major morphological and geometric model parameters. 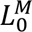, optimal muscle fascicle length; 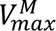/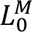, normalized maximum fascicle shortening velocity; SO, percent of slow-twitch muscle fibers; PCSA, physiological cross-sectional area; PA, pennation angle; *a*_1_ and *φ*_1_, parameters defining the origin of MTU (see Fig. 2 and Table 7); 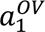 and 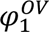, parameters defining the origin of MTU path between the origin and the via point; *a*_2_ and *φ*_2_, parameters defining the insertion of MTU; 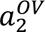 and 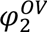, parameters defining the insertion of MTU path between the origin and the via point; 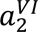 and 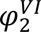, parameters defining the insertion of MTU path between the via point and insertion; R_1_ and R_2_, radii of joints spanned by a two-joint MTU; The maximal moments of force were computed from all studied cats (n=7) and walking cycles (n=211), see Table 2.

We determined MTU activations during walking by minimizing the function of muscle fatigue (Crowninshield and Brand, 1981). This and similar optimization criteria minimizing the sense of fatigue and effort, metabolic energy expenditure, and movement variability have demonstrated reasonable performance predicting recorded muscle activations or muscle forces during human and cat locomotion and other skilled/reflex motor behaviors (Crowninshield and Brand, 1981; Prilutsky et al., 1997; Prilutsky, 2000b, a; Anderson and Pandy, 2001; Haruno and Wolpert, 2005). Comparing the activation patterns of 7 forelimb muscles computed from the 40-MTU model with the corresponding EMG recordings during walking in this study, we can see that only 5 MTUs demonstrate some qualitative agreement (Fig. 8; SPS, TLAT, TLONG, PT, BR). Note that we cannot compare the magnitudes of the computed and recorded activity (the EMG magnitude is arbitrary). We also need to account for the muscle activation and deactivation times that are ignored in our static optimization; this partially explains the earlier EMG onset and offset compared to computed activations in some MTUs (e.g. SPS, TLAT, TLONG, PT, BR) but are accounted for in dynamic optimization (Anderson and Pandy, 2001; Neptune et al., 2008; Prilutsky et al., 2016). The earlier appearance of muscle EMG compared to MTU force production depends on the electromechanical delay caused in part by developing tendon tension, which is muscle- and task-dependent. Muscles with shorter tendons undergoing lengthening actions tend to have shorter electromechanical delays (Komi, 2003). In general, computed forelimb activations less accurately matched recorded muscle activity or force compared to the cat hindlimb (Prilutsky et al., 1997) or human leg muscles (Crowninshield and Brand, 1981; Anderson and Pandy, 2001). This can be explained by several factors, including a more complex organization of forelimb/arm muscle morphology, with a greater number of muscle compartments with separate innervations and different activity during a single motor task (English and Weeks, 1987; Brown et al., 2007). Thus, our computed MTU activations should be considered with caution. Nevertheless, the main activity bursts of major extensor and flexor MTUs were reproduced by computed activations of the 40-MTU model (SPS, TLAT, TLONG, PT, BR) with only partial reproduction of EMG patterns of two-joint muscles (BB, ECR) (Fig. 8).

We also assumed that the proportion of proprioceptive axons entering a spinal cord segment from a given MTU corresponds to the proportion of motoneurons of this MTU in the same segment. Although, in general, somatotopy between muscle motoneurons and corresponding muscle afferents has been established (Sterling and Kuypers, 1967b, a; Fritz et al., 1986a; Fritz et al., 1986b; Levine et al., 2012), we do not know if proportions of motoneurons and proprioceptive afferents entering the same spinal segments are the same. We found only one study in 2 monkeys in which the number of motoneurons and dorsal root ganglion neurons labeled by the retrograde transport of horseradish peroxidase from the same muscle (EDC) were determined in segments C5-T2 and showed only rough qualitative agreement (Jenny and Inukai, 1983).

### Functional significance of cat forelimb morphology

#### Morphology comparison between cat forelimbs and hindlimbs

Morphological data of the cat forelimb were obtained to better understand the role of forelimb muscles in generating motor output and sensory feedback during locomotion. A comparison of cat forelimb morphological characteristics with corresponding data of the hindlimb (Sacks and Roy, 1982) revealed a number of similar and contrasting features that support the functional specializations of the two limbs. Muscle mass of the cat forelimb (Fig. 3A, Table 5) and hindlimb (Sacks and Roy, 1982) is concentrated proximally, which contributes to reducing the limb moment of inertia with respect to the proximal joints and energy expenditure required to transport the leg during the swing phase (see Introduction). PCSAs also decline in a proximal-distal direction in the forelimb (Fig. 3B, Table 5), but are more equally distributed in the hindlimb (Sacks and Roy, 1982). The total mass and PCSA of the shoulder protractors (anatomically analogous to hip flexors) is 3-4 times greater than mass and PCSA of shoulder retractors (Fig. 3A, B), although this imbalance would probably be lessened by inclusion of extrinsic shoulder muscles like latissimus dorsi and pectoralis major. The hip flexors in the cat are smaller and weaker than the hip extensors (Sacks and Roy, 1982). The discrepancy in PCSA and muscle mass, which is proportional to muscle volume or the product of mean PCSA and muscle fascicle length, can be explained by the unique role of the shoulder protractors in supporting body weight in stance and absorbing mechanical energy of the body in the walking cycle at constant speed. This is reflected in a greater area of the negative force impulse of the horizontal ground reaction force F_x_ compared to the positive one (Fig. 4F). Note that muscle volume indicates the maximum amount of positive work (or generated energy) or negative work (absorbed energy) the muscle can do because PCSA is proportional to the muscle’s maximum isometric force (Powell et al., 1984), and fascicle length is proportional to the length range of active force production. Thus, shoulder protractors are well suited to absorb energy of the body during stance when these muscles produce the resultant protraction moment of force at the shoulder (Fig. 4G; see also Fig. 7A, B, C) and are increasing in length for the first two-thirds of stance (Fig. 5A, B). Elbow extensors and wrist plantarflexors do not contribute substantially to energy absorption, or body deceleration, during stance because there is a very limited joint angle yield in early stance at these joints (Fig. 4B, C) and corresponding lengthening of these muscles (Fig. 5A, B). The analogous knee and ankle joints of the cat hindlimb have much more profound joint yield and energy absorption in early stance (McFadyen et al., 1999; Prilutsky et al., 2011).

In addition, the total mass of forelimb muscles was substantially smaller than that of the hindlimb, i.e. 139.8 g, Table 5 vs 231.7 g, Table 1 in (Sacks and Roy, 1982), respectively, and thus can contribute less to energy generation for propulsion in locomotion and jumping. The shoulder adductor, abductor and protractor muscles have substantial force generating capacities (large PCSA) compared to the retractors (Fig. 3B). Hindlimb adductors, in contrast, have substantially smaller total PCSA than hip extensors. These differences may reflect the requirement for rapid non-sagittal force production and movements of the forelimbs for lateral balance control in locomotion (Park et al., 2019) and landing (McKinley et al., 1983), as well as defensive fighting or prey capture.

We found considerable variability in fascicle length between forelimb functional MTU groups, e.g., flexors and extensors vs. adductors, supinators, and pronators (Fig. 3C) and within groups, e.g. brachioradialis (BCD, 89.6 mm) vs. brachialis (BR, 32.4 mm) for elbow flexors (Table 5). Similar variability was observed in the cat hindlimb, e.g. soleus (41.7 mm) vs. plantaris (18.7 mm) for ankle extensors (Sacks and Roy, 1982). The within-group variation of fascicle length can support functional diversity. One special case, BCD, an elbow flexor and forearm supinator with low force producing potential (PCSA = 0.08 cm^2^, more than an order of magnitude smaller than in its synergists, Table 5) has substantially longer fascicles than major elbow flexors or supinators. The BCD may bear some similarity to the abductor cruris caudalis or tenuissimus in the hindlimb (Lev-Tov et al., 1988). Both muscles span several joints/DOFs and their small PCSA indicates that they produce little force. The sensory functions of these muscles may be more important, as they could signal whole limb length (note the pattern similarity between the forelimb length (Fig. 4D) and the BCD MTU length (Fig. 5A) and estimated spindle group II activity (Fig. 9B) during the walking cycle).

#### Role of the forelimb in weight support and postural control

Weight support during quiet standing requires relatively low force production over a long period with little postural change. This may be particularly true for the cat forelimb, which remains rigid and extended during perturbations of quiet standing (Lacquaniti and Maioli, 1994) and may transfer substantial load through bony structures. The forelimb and hindlimb both possess muscles with predominantly slow-twitch fiber types. The ankle extensor soleus is composed of 100% slow fibers (Ariano et al., 1973), and the ANC and TMED contain 100% and 83% slow-twitch fibers, respectively (Collatos et al., 1977). These antigravity muscles are therefore mechanically and metabolically suited for postural tasks. During horizontal perturbations of the support surface, the hindlimbs exhibit anisotropic force responses in which the greatest stiffness lies backward and outward, while the forelimbs exhibit similar magnitudes of force response in all directions (Macpherson, 1988; Honeycutt and Nichols, 2010). The greater isotropy in force production potential among the shoulder protractors, adductors and abductors (Fig. 3B) compared to corresponding muscles at the hip may explain at least in part the more radially symmetric force responses of the forelimb to postural perturbations. This supports the idea that the forelimbs function primarily as struts during postural control (Rushmer et al., 1983; Lacquaniti and Maioli, 1994).

#### Role of the forelimb in locomotion

The requirements for locomotion include weight-support, propulsion, balance control, and mechanisms for rapid change of direction. Previous work contrasted the forelimbs of the hare (Williams et al., 2007) and cheetah (Hudson et al., 2011a), specialized for rapid changes in direction and speed, with the greyhound (Williams et al., 2008), specialized for straight-line sprinting. The forelimb of the cat, like the forelimb of the hare and cheetah, appears especially well suited for weight-support, stability control, and turning. Unlike the greyhound, which has nearly equal distribution of muscle mass between the forelimbs and hindlimbs (the forelimb muscles are only 11% lighter), the mass of the forelimb muscles in the cat is 60% of the hindlimb muscles – our Table 5 and Table 1 in (Sacks and Roy, 1982). This suggests much lower total power generating capacity of the forelimbs and a limited contribution to propulsion. The cat, like the hare (Williams et al., 2007) and the cheetah (Hudson et al., 2011b, a), displays substantial isotropy for PCSA among the protractor, adductors and abductors at the shoulder (Fig. 3B and Table 5). This lack of specialization for protraction-retraction suggests that forelimb adductors and abductors, which are co-active during stance (English, 1978a, b), can produce relatively greater mediolateral forces, providing stability and control of direction during stance. The relatively large PCSA of shoulder adductors and abductors also contributes to the larger moment arms of these muscles with respect to the corresponding axis of rotation at the shoulder, given that the centroid path of the muscle passes further away from this axis. This in turn leads to greater opposing adduction and abduction moments of force at the shoulder, contributing to lateral stabilization of the forelimb during the stance phase of locomotion. The relative similarity in average fascicle length between protractors-retractors and extensors-flexors at the shoulder and elbow in the cat (Fig. 3C, Table 5) also suggests a lesser requirement for acceleration and propulsion in the forelimb than in the hindlimb, where the powerful quadriceps and triceps surae have fascicles half the length of their antagonists (Sacks and Roy, 1982). Therefore, the above characteristics of the cat forelimb musculature appears to be better suited for lateral stability control and for changing the direction of locomotion than for producing acceleration and power for locomotion.

#### Role of the forelimb in manipulation

Reaching tasks follow a consistent sequence of elbow flexion and wrist dorsiflexion with shoulder protraction as the cat lifts the limb, followed by elbow extension, as the cat thrusts the limb forward, and supination of the antebrachium and wrist plantarflexion prior to target contact; in the retrieval stage, the wrist dorsiflexes and the elbow flexes, while the limb pronates (Illert, 1996; Yakovenko and Drew, 2015). Of special note is the presence of powerful supination and pronation of the forearm about its long axis, the function that is essentially lacking in the homologous hindlimb segment, the shank. In the distal forelimb, the supinator (SUP) and PT are highly specialized, low-mass muscles with short, highly pennated fascicles (Table 5) that provide high forces needed for prey retrieval and suppression, respectively. In the hindlimb, antigravity muscles have much shorter fascicles than their antagonists, often described as specialization for force production (Lieber and Ward, 2011). This specialization is muted in the forelimb, possibly reflecting the benefit of powerful flexors for prey capture as a counter to the benefit of powerful extensors during pursuit. Non-sagittal movements are also frequently involved in foraging and exploratory movements and prey capture. The relatively larger masses of shoulder abductors and adductors are well suited to these out-of-sagittal plane movements and provide a general-purpose design for strut-like weight support, controlling balance and direction during locomotion and reaching in many directions.

#### Contribution of cat forelimb morphology to somatosensory control of locomotion

Motion-dependent proprioceptive input to spinal locomotor networks depends on muscle fascicle length, velocity of stretch, and force produced by the MTU (Prochazka, 1999). Several morphological and geometric characteristics of MTUs affect the above mechanical variables. A short muscle fascicle length with respect to tendon length can substantially reduce both the range of length change and stretch velocity of the fascicles during locomotion. This is because the relatively long and elastic tendon takes up a substantial amount of stretch in an active MTU, as observed in the cat’s medial gastrocnemius, with short fascicles, but not in its close synergist, the soleus, with much longer fascicles, during the stance phase of level walking (Maas et al., 2009). In the cat hindlimb, proximal MTUs (two-joint hamstrings, sartorius) have much longer muscle fascicles/belly than distal MTUs (triceps surae) (Sacks and Roy, 1982; Scott et al., 1992). These morphological differences support distinct roles of length-dependent sensory input from proximal hip muscles in regulating phase transitions during cat locomotion (Kriellaars et al., 1994; Lam and Pearson, 2002; McVea et al., 2005; Gregor et al., 2006). In the cat forelimb, although on average fascicle length of proximal MTUs is not markedly different from that of more distal MTUs (Fig. 3D), one-joint shoulder retractors (SD, TN) and the two-joint shoulder protractor-elbow flexor BB have relatively long fascicle lengths (25.1 mm – 46.1 mm, Table 5). The MTU length (Fig. 5A) and estimated activity of muscle spindle group II afferents (Fig. 9B) reach peak values at the end of swing and stance, respectively. Thus, the length-dependent sensory feedback from these MTUs are strong candidates for communicating to rhythm generating spinal networks that the forelimb is approaching phase transitions. Properties of spindle group II afferents are well suited to sense limb position (Banks et al., 2021). The BB’s patterns of MTU length (Fig. 5A) and spindle group II activity (Fig. 9B) closely match the pattern of forelimb orientation (Fig. 4E) during the walking cycle. Additionally, lengthening patterns of muscle fascicles and spindles in shoulder retractors during swing and in BB during stance are not likely to be distorted by their active shortening as these MTUs are not active during the corresponding phases of the walking cycle (English, 1978a, b; Krouchev et al., 2006), see also Fig. 8A. Two slightly more distant MTUs, the elbow flexor-forearm supinator BCD and the elbow flexor-wrist dorsiflexor ECR have very long fascicle lengths (89.6 mm and 47.3 mm, Table 5). Their MTU length patterns (Fig. 5A) and estimated spindle group II activity (Fig. 9B) closely match forelimb length changes during the walking cycle (Fig. 4D). Thus, length-dependent inputs from these two MTUs could provide the nervous system with information about overall limb length.

The forelimb moment arms are greater for shoulder protractors and retractors than for more distal muscles as evident from the muscle path locations with respect to the joint centers (Figs 1A and 5C). This increases MTU length changes for a given change in joint angle, making the muscle spindles in the proximal muscles more sensitive to joint position and motion (Hall and McCloskey, 1983). Thus, the relatively long moment arms and fascicle lengths are morphological characteristics of proximal forelimb muscles that make them well suited for providing limb orientation and angular motion information to the spinal cord.

The limb posture also influences the precision of limb position sense and endpoint position control. The limb posture constrains, to a large extent, the limb endpoint precision and stiffness ellipses, which are perpendicular to each other (Bunderson et al., 2010; Burkholder, 2016; Oh and Prilutsky, 2022).

Group Ib Golgi tendon afferents from ankle extensors transmit load-dependent information to locomotor networks (Duysens and Pearson, 1980; Guertin et al., 1995; Hiebert and Pearson, 1999; Ross and Nichols, 2009). This information affects the timing of the stance-swing transition and regulates the activity of hindlimb extensors. It is currently unknown whether group Ib afferents of forelimb distal extensors produce similar functional effects, although they are likely discharging during stance (Fig. 9C) as these antigravity muscles are active and produce moments of force (Figs. 4I, 7A-C).

### Future directions

The results of this study are essential for understanding the role of cat forelimb morphology for locomotor sensorimotor functions and in providing important information about the potential role of somatosensory feedback in the neural control of quadrupedal locomotion. Although previous studies established strong effects of forelimb movements and forelimb sensory stimulation on hindlimb movements and reflex responses (Miller et al., 1973; Schomburg et al., 1998; Doperalski et al., 2011; Frigon, 2017; Hurteau et al., 2018; Merlet et al., 2022; Harnie et al., 2024; Mari et al., 2024), this information has not been incorporated into anatomically and neurophysiologically accurate models of mammalian quadrupedal locomotion. These types of models should be capable of providing mechanistic interpretations and explanations of a wide range of experimental data on mechanics and muscle activity patterns during quadrupedal locomotion, including those resulting from various lesions in the spinal cord (Doperalski et al., 2011; Audet et al., 2023; Danner et al., 2023; Shepard et al., 2023; Mari et al., 2024). Spinal cord injury in people, especially complete injury, remains a major healthcare problem. Translational studies on relatively large animal models, like cats (Gerasimenko et al., 2009; Frigon, 2020), and their detailed neuromechanical models (Markin et al., 2016; Nichols et al., 2016) are indispensable for understanding spinal and somatosensory control of locomotion and for developing new rehabilitation approaches. One new promising rehabilitation method is spinal cord stimulation through implanted epidural or noninvasive transcutaneous dorsal stimulating electrodes that appears to increase excitability of spinal locomotor networks transynaptically by activating somatosensory afferents entering the dorsal horns under the electrodes (Harkema et al., 2011; Minassian et al., 2016; Lorach et al., 2023). The spatiotemporal maps of sensory inputs to cervical spinal segments obtained in this study (Fig. 10B-D) could guide the development of targeted epidural/transcutaneous stimulation to improve locomotor function after injury because cervical stimulation has a strong impact on locomotor activity of leg muscles in humans (Gerasimenko et al., 2015). We plan to incorporate the forelimb musculoskeletal model with that of the hindlimbs (Markin et al., 2016) to develop a neuromechanical model of quadrupedal locomotor control to address the above goals.

## Supporting information

Supplemental Table 1_3D forelimb morphology

Supplemental Table 2_2D forelimb morphology

Supplement 3_Description of muscle wrapping

## Author contributions

B.I.P., T.J.B., A.F and I.A.R. conceived and designed research. R.S.M., N.E.B., T.R.N. and T.J.B. performed forelimb dissections and morphological measurements and modeling. A.N.K. and B.I.P. recorded mechanics of locomotion. A.F. provided EMG recordings. S.M.R. developed the 2D forelimb model and computed motor and sensory signals during locomotion. A.N.K. and B.I.P. developed code for and performed inverse dynamics and EMG analyses. S.M.R. and J.A.M. performed sensitivity analysis of the model. S.M.R. and B.I.P. prepared figures. S.M.R., R.S.M., N.E.B, T.J.B. and B.I.P. drafted manuscript. All authors discussed, reviewed, edited, and approved the final version of the manuscript.

## Data availability

Raw morphological data are provided in supplemental material. Additional data will be made available on request.

MATLAB code for computing all variables presented in the article is available at https://github.com/smarahmati/CatForelimbNeuromechanics.

## Supplementary data

Supplement 1: 3D coordinates of forelimb muscle origins, insertions, and joint centers

Supplement 2: 2D coordinates of forelimb muscle origins, insertions, and joint centers

Supplement 3: Equations describing muscle wrapping around joints.

