## Supplement 3_Description of muscle wrapping for "ROLE OF FORELIMB MORPHOLOGY IN MUSCLE SENSORIMOTOR FUNCTIONS DURING LOCOMOTION IN THE CAT"

### Determining muscle-tendon unit length and moment arms of cat forelimb muscles

#### Overview

We developed equations to calculate muscle-tendon unit (MTU) lengths as a function of joint angles in the sagittal plane. Then we computed the MTU moment arm at a joint as the derivative of MTU length over the joint angle. We determined MTU lengths parametrically for a one-joint muscle, a two-joint muscle and a muscle with via points. We introduced a criterion to determine if a MTU wraps around a joint. We considered the joint surface to be a circle. If the MTU wraps around a joint, the wrapping length was included in computations of MTU lengths, and the moment arm was equal to the joint radius.

#### Geometric model

We introduced parameters $f_i$ and $n_i$ ($i = 1$ for one-joint MTU and $i = 1,2$ for two-joint MTU) that define location of the MTU path with respect to the joint centers. If MTU passes posteriorly of the joint, $f_i = 1$; if it passes anteriorly, $f_i = -1$. If the joint angle is defined on the posterior side, $n_i = 1$; if it is defined on the anterior side, $n_i = -1$. Using these parameters and mathematical modeling, we introduced a criterion ($\theta_i$) to determine whether the MTU wraps around a joint ($\theta_i \geq 0$) or not ($\theta_i < 0$).

##### *Length computation of one-joint MTU as a function of joint angle*

We considered four possible cases for one-joint MTU path with respect to the joint. These cases are categorized by posterior-anterior location of MTU path with respect to the join and by whether the MTU path is in contact with joint surface (Fig. S1). MTU either wraps around the joint ($\theta \geq 0$) or there is no contact between the MTU and joint surface ($\theta < 0$). The parameter $\theta_1$ is computed as follows (see Fig. S1):

$$\gamma_1 = f_1 \arccos\left(\frac{R_1}{a_1}\right) \quad \text{(S1)}$$

$$\gamma_2 = -f_1 \arccos\left(\frac{R_1}{a_2}\right) \quad \text{(S2)}$$

$$\beta_1 = \gamma_1 + \varphi_1 \quad \text{(S3)}$$

$$\beta_2 = \gamma_2 + \varphi_2 \quad \text{(S4)}$$

$$\theta_1 = |f_1 - n_1|\pi + f_1 n_1(\alpha_1 - n_1(\beta_1 - \beta_2)), \quad \text{(S5)}$$

where $0 \leq \alpha \leq 2\pi$ is the joint angle, $a_1$ and $a_2$ are distances between the joint center and MTU origin and insertion, respectively; $\varphi_1$ is the angle between the line connecting MTU origin with joint center and the proximal segment; $\varphi_2$ is the angle between the line connecting MTU insertion with joint center and the distal segment; and $R_1$ is joint radius.

With a known value of parameter $\theta_1$, MTU length is computed for each case in Fig. S1 as follows.

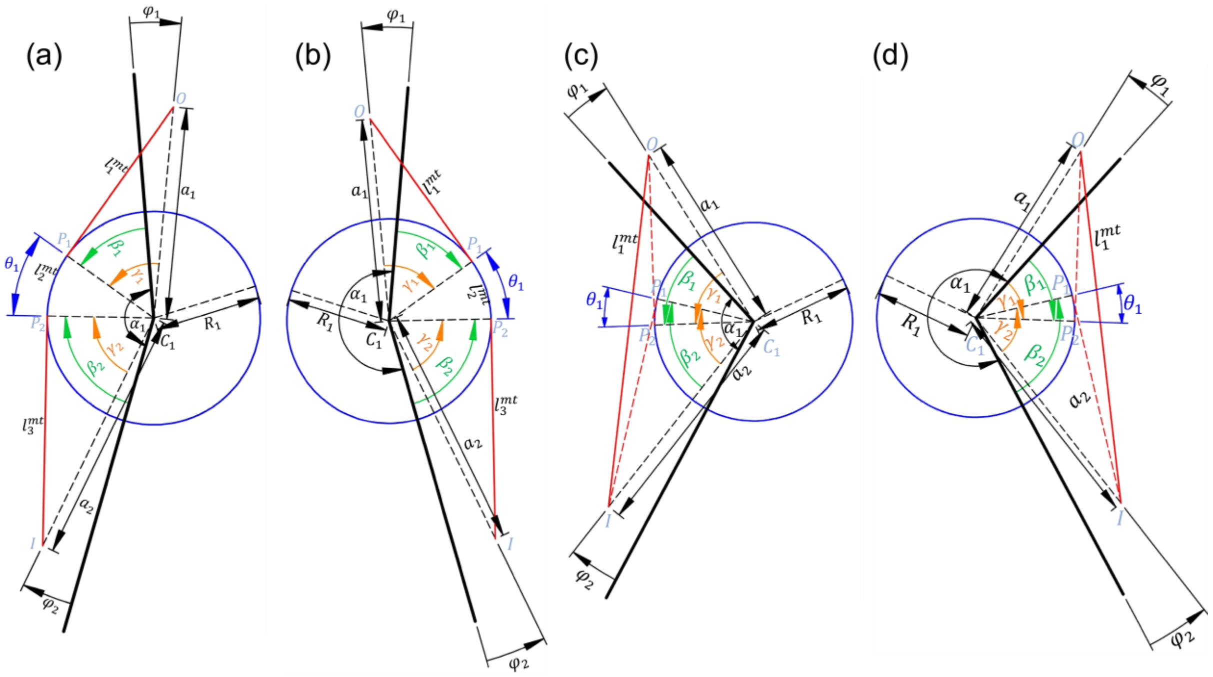

**Figure S1**. Possible cases of one-joint MTU path location with respect to the joint. In all cases $n_1 = 1$. (a) MTU wraps joint surface and is located posterior to the joint center ($f_1 = 1$). (b) MTU wraps joint surface and is located anterior to the joint ($f_1 = -1$). (c) MTU is not in contact with joint surface and is located posterior to the joint ($f_1 = 1$). (d) MTU is not in contact with joint surface and is located anterior to the joint ($f_1 = -1$).

*Cases (a) and (b) where* $(\theta_1 \geq 0)$

$$l_1^{mtu} = \sqrt{a_1^2 - R_1^2} \tag{S6}$$

$$l_2^{mtu} = R_1\theta_1 \tag{S7}$$

$$l_3^{mtu} = \sqrt{a_2^2 - R_1^2} \tag{S8}$$

$$l^{mtu} = l_1^{mtu} + l_2^{mtu} + l_3^{mtu} \tag{S9}$$

*Cases (c) and (d) where* $(\theta_1 < 0)$

$$l_1^{mtu} = \sqrt{a_1^2 + a_2^2 - 2a_1a_2\cos\left(\alpha_1 - n_1(\varphi_1 - \varphi_2)\right)}, \tag{S10}$$

$$l^{mtu} = l_1^{mtu} \tag{S11}$$

where $l^{mtu}$ is the total MTU length.

#### *Computation of two-joint muscle length as a function of joint angles*

We consider 16 possible cases of a two-joint MTU path categorized by wrapping around proximal and/or distal joint and by the posterior-anterior path location with respect to the two joints. Figures S2-S5 show all the cases. The same criterion for wrapping ($\theta_i \geq 0$) or no contact with joint surface ($\theta_i < 0$) was used for each joint. Parameter $\theta_i$, $i = 1,2$ is computed as follows (see Figs. S2-S5):

$$\gamma_1 = f_1 \arccos\left(\frac{R_1}{a_1}\right) \tag{S12}$$

$$\gamma_2 = -f_2 \arccos\left(\frac{R_2}{a_2}\right) \tag{S13}$$

$$\beta_1 = \gamma_1 + \varphi_1 \tag{S14}$$

$$\beta_2 = \gamma_2 + \varphi_2 \tag{S15}$$

$$h_1 = \sqrt{l_s^2 - (f_1 R_1 - f_2 R_2)^2} \tag{S16}$$

$$h_2 = \sqrt{R_2^2 + h_1^2} \tag{S17}$$

$$h_3 = \sqrt{R_1^2 + h_1^2} \tag{S18}$$

$$\rho_1 = -f_1 \arccos\left(\frac{h_2^2 - R_1^2 - l_s^2}{-2R_1 l_s}\right) \tag{S19}$$

$$\rho_2 = f_2 \arccos\left(\frac{h_3^2 - R_2^2 - l_s^2}{-2R_2 l_s}\right) \tag{S20}$$

$$\theta_i = |f_i - n_i|\pi + f_i n_i(\alpha_i + (-1)^i n_i(\beta_i - \rho_i)), \tag{S21}$$

where $i = 1{,}2$; $l_s$ is segment length; $R_2$ is the radius of the proximal joint. Equations S12-S21 are valid for all sixteen cases in Figs. S2-S5. The parameters $\theta_1$ and $\theta_2$ are defined for the proximal and distal joint, respectively. Two-joint MTU length is computed for different cases as follows:

*Cases where* $(\theta_1 \geq 0, \theta_2 \geq 0)$; Fig. S2

$$l_1^{mtu} = \sqrt{a_1^2 - R_1^2} \tag{S22}$$

$$l_2^{mtu} = R_1\theta_1 \tag{S23}$$

$$l_3^{mtu} = h_1 \tag{S24}$$

$$l_4^{mtu} = R_2\theta_2 \tag{S25}$$

$$l_5^{mtu} = \sqrt{a_2^2 - R_2^2} \tag{S26}$$

$$l^{mtu} = l_1^{mtu} + l_2^{mtu} + l_3^{mtu} + l_4^{mtu} + l_5^{mtu} \tag{S27}$$

*Cases where* $(\theta_1 \leq 0, \theta_2 \geq 0)$; Fig. S3

$$h_4 = \sqrt{l_s^2 + a_1^2 - 2a_1 l_s \cos(\alpha_1 - n_1\varphi_1)} \tag{S28}$$

$$l_1^{mtu} = \sqrt{h_4^2 - R_2^2} \tag{S29}$$

$$\rho_2^{m_1} = f_1 arcos\left(\frac{a_1^2 - l_s^2 - h_4^2}{-2l_s h_4}\right) \tag{S30}$$

$$\rho_2^{m_2} = f_2 arcos\left(\frac{R_2}{h_4}\right) \tag{S31}$$

$$\rho_2^m = f_2\left|\rho_2^{m_1} + \rho_2^{m_2}\right| \tag{S32}$$

$$\theta_2^m = |f_2 - n_2|\pi + f_2 n_2(\alpha_2 - n_2(\beta_2 - \rho_2^m)) \tag{S33}$$

$$l_2^{mtu} = R_2\theta_2^m \tag{S34}$$

$$l_3^{mtu} = \sqrt{a_2^2 - R_2^2} \tag{S35}$$

$$l^{mtu} = l_1^{mtu} + l_2^{mtu} + l_3^{mtu} \tag{S36}$$

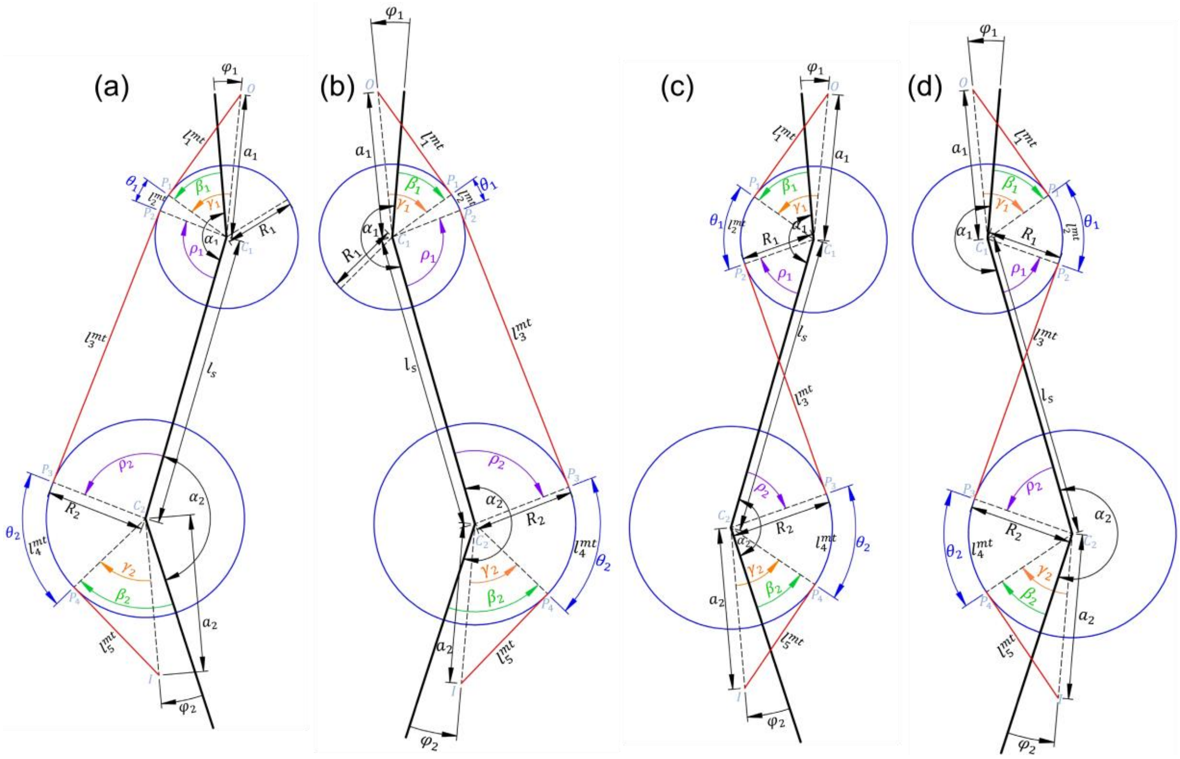


**Figure S2**. Possible cases of two-joint MTUs for which $\theta_1 \geq 0, \theta_2 \geq 0$. For all 4 cases the values of $n_1$ and $n_2$ are the same ($n_1 = 1, n_2 = -1$). (a) The MTU is located posterior to both proximal and distal joints ($f_1 = 1, f_2 = 1$). (b) The MTU is located anterior to both proximal and distal joints ($f_1 = -1, f_2 = -1$). (c) The MTU is located posterior to the proximal joint and anterior to the distal joint ($f_1 = 1, f_2 = -1$). (d) The MTU is located anterior to the proximal joint and posterior to the distal joint ($f_1 = -1, f_2 = 1$).

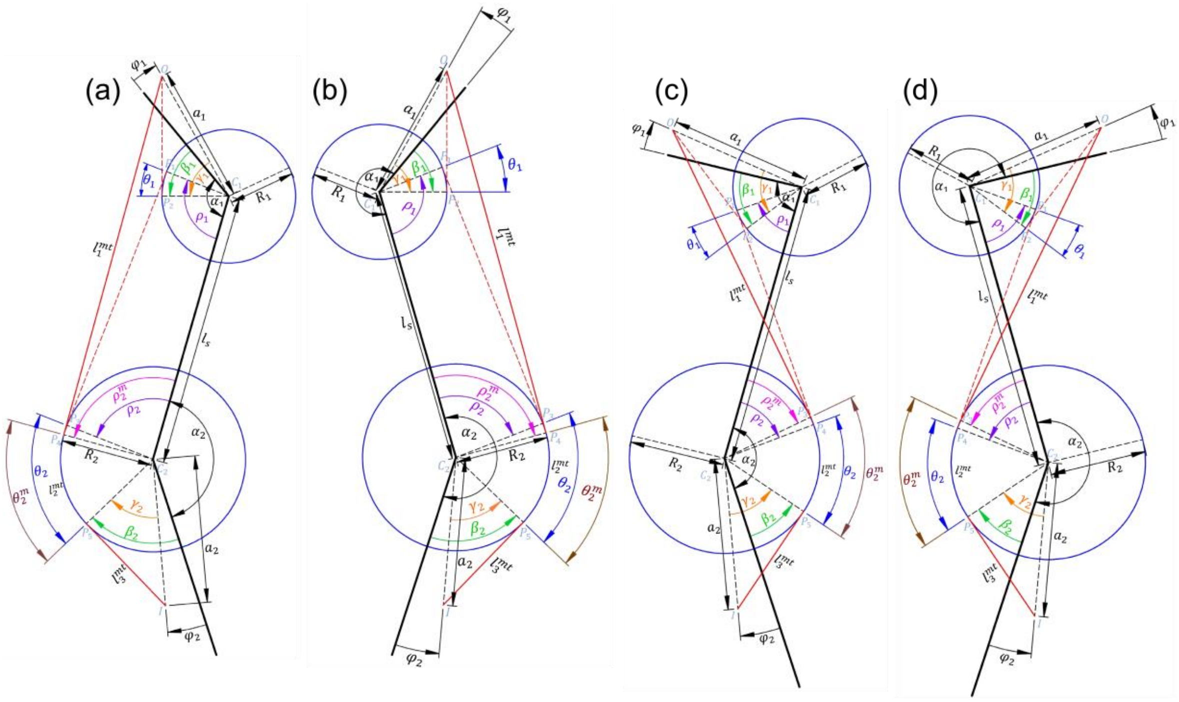

**Figure S3**. Possible cases of two-joint MTUs for which $\theta_1 \leq 0, \theta_2 \geq 0$. For all 4 cases the values of $n_1$ and $n_2$ are the same ($n_1 = 1, n_2 = -1$). (a) The MTU is located posterior to both proximal and distal joints ($f_1 = 1, f_2 = 1$). (b) The MTU is located anterior to both proximal and distal joints ($f_1 = -1, f_2 = -1$). (c) The MTU is located posterior to the proximal joint and anterior to the distal joint ($f_1 = 1, f_2 = -1$). (d) The MTU is located anterior to the proximal joint and posterior to the distal joint ($f_1 = -1, f_2 = 1$).

*Cases where* ($\theta_1 \geq 0, \theta_2 \leq 0$); Fig. S4

$$l_1^{mtu} = \sqrt{a_1^2 - R_1^2} \tag{S37}$$

$$h_5 = \sqrt{l_s^2 + a_2^2 - 2a_2 l_s \cos\left(\alpha_2 + n_2\varphi_2\right)} \tag{S38}$$

$$\rho_1^{m_1} = f_1 arcos\left(\frac{a_2^2 - l_s^2 - h_5^2}{-2l_s h_5}\right) \tag{S39}$$

$$\rho_1^{m_2} = f_2 arcos\left(\frac{R_2}{h_5}\right) \tag{S40}$$

$$\rho_1^m = -f_1\left|\rho_1^{m_1} + \rho_1^{m_2}\right| \tag{S41}$$

$$\theta_1^m = |f_1 - n_1|\pi + f_1 n_1(\alpha_1 - n_1(\beta_1 - \rho_1^m)) \tag{S42}$$

$$l_2^{mtu} = R_1\theta_1^m \tag{S43}$$

$$l_3^{mtu} = \sqrt{h_5^2 - R_1^2} \tag{S44}$$

$$l^{mtu} = l_1^{mtu} + l_2^{mtu} + l_3^{mtu} \tag{S45}$$

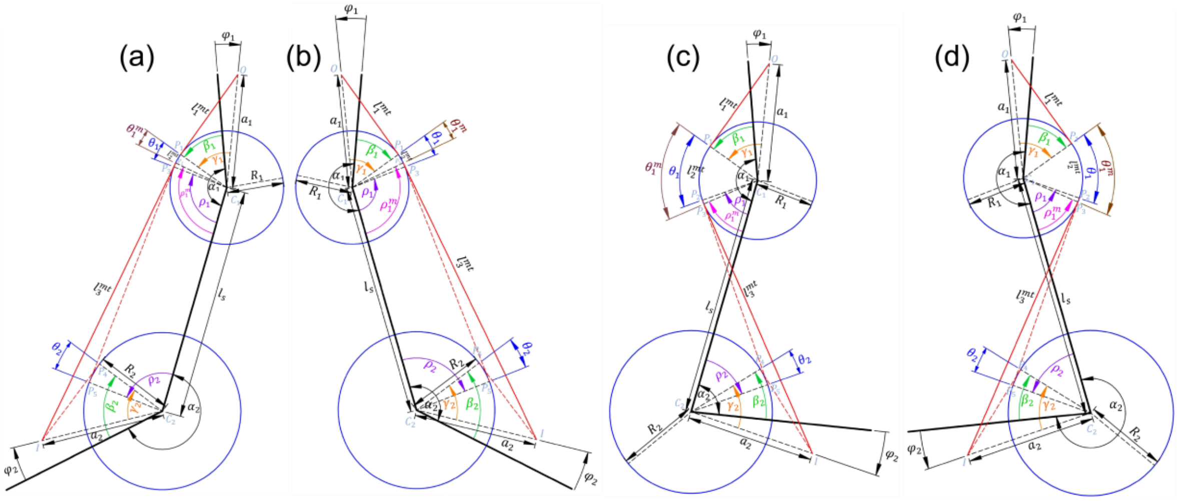

**Figure S4**. Possible cases of two-joint MTUs for which $\theta_1 \geq 0, \theta_2 \leq 0$. For all 4 cases the values of $n_1$ and $n_2$ are the same ($n_1 = 1, n_2 = -1$). (a) The MTU is located posterior to both proximal and distal joints ($f_1 = 1, f_2 = 1$). (b) The MTU is located anterior to both proximal and distal joints ($f_1 = -1, f_2 = -1$). (c) The MTU is located posterior to the proximal joint and anterior to the distal joint ($f_1 = 1, f_2 = -1$). (d) The MTU is located anterior to the proximal joint and posterior to the distal joint ($f_1 = -1, f_2 = 1$).

*Cases where* ($\theta_1 \leq 0, \theta_2 \leq 0$); Fig. S5

$$h_6 = a_1 \cos(\pi - \alpha_1 + n_1\varphi_1) \tag{S46}$$

$$h_7 = a_2 \cos(\pi - \alpha_2 - n_2\varphi_2) \tag{S47}$$

$$h_8 = l_s + h_6 + h_7 \tag{S48}$$

$$l_1^{mtu} = \sqrt{(f_1\sqrt{a_1^2 - h_6^2} - f_2\sqrt{a_2^2 - h_7^2})^2 + h_8^2} \tag{S49}$$

$$l^{mtu} = l_1^{mtu} \tag{S50}$$

where $l^{mtu}$ is the total MTU length for all cases.

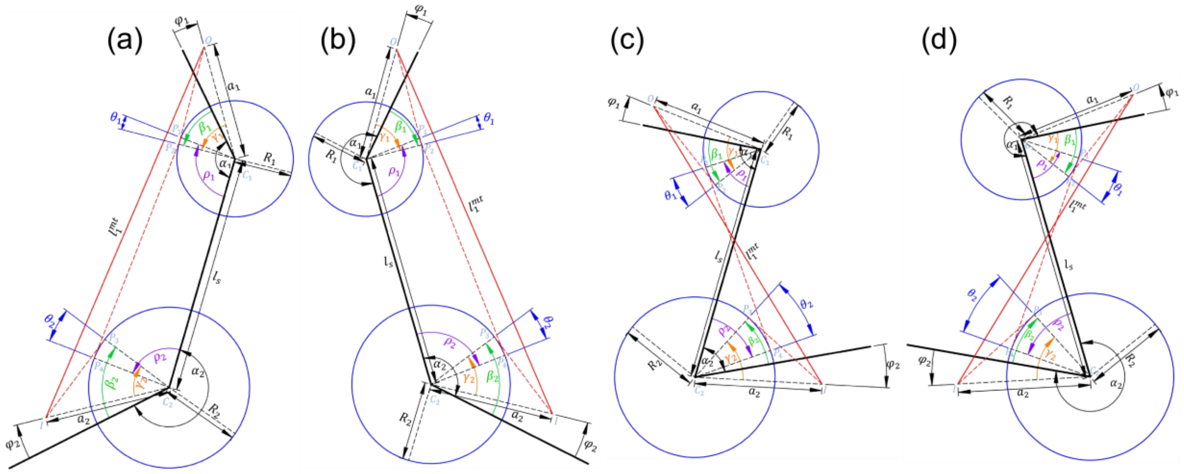

**Figure S5**. Possible cases of two-joint MTUs for which $\theta_1 \leq 0, \theta_2 \leq 0$. For all 4 cases the values of $n_1$ and $n_2$ are the same ($n_1 = 1, n_2 = -1$). (a) The MTU is located posterior to both proximal and distal joints ($f_1 = 1, f_2 = 1$). (b) The MTU is located anterior to both proximal and distal joints ($f_1 = -1, f_2 = -1$). (c) The MTU is located posterior to the proximal joint and anterior to the distal joint ($f_1 = 1, f_2 = -1$). (d) The MTU is located anterior to the proximal joint and posterior to the distal joint ($f_1 = -1, f_2 = 1$).

***Computation of MTU length with via points as a function of joint angles***

A muscle with via points can be simply divided into several parts including the origin to the first via point, first via point to the second via point and so on to the insertion. Each part can be considered as a one-joint MTU, two-joint MTU or MTU with a constant length, i.e. a MTU does not span a joint. The length of each MTU part can be computed based on the above equations. Therefore, the total MTU length is the sum of each part length.

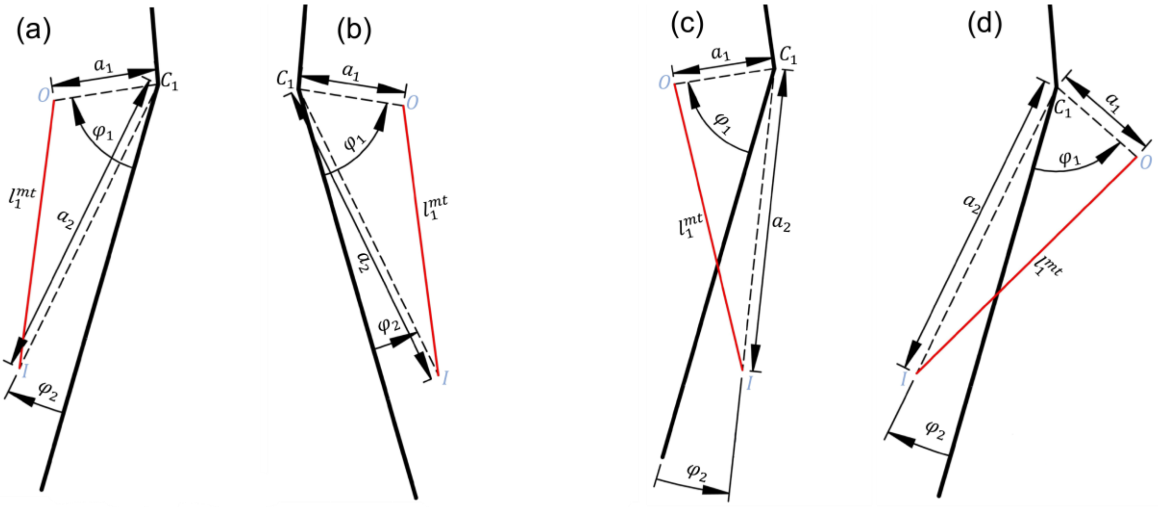

**Figure S6**. Different cases of MTU with constant length. (a) The muscle is located on posterior side of the segment. (b) The muscle is located on anterior interior side of the segment. (c) The MTU origin is located on posterior side of the segment while the attachment, on anterior side. (d) The MTU origin is located anterior side of the segment while the attachment, on posterior side.

There is a special case in which muscle attachment points are on the same segment; therefore, the length of this MTU part is constant and does not depend on a joint angle (Fig. S6). The length of this part can be computed based on MTU attachment parameters:

$$l_1^{mtu} = \sqrt{a_1^2 + a_2^2 - 2a_1 a_2 \cos(\varphi_1 - \varphi_2)}. \quad \text{(S51)}$$

Based on the above formulations of path length for one-joint, two-joint MTUs and MTUs with constant length, the total MTU length can be parametrically computed as a function of joint angles and then differentiated with respect to each joint angle to obtain MTU moment arms.
